# Synergistic effects of deleting the tyrosine phosphatases Shp1 and Shp2 on megakaryopoiesis and thrombopoiesis in mice

**DOI:** 10.1101/2025.10.24.684367

**Authors:** Elsa Barré, Marc-Damien Lourenco-Rodrigues, Lucie Zimmermann, Marion Pugliano, Cécile Loubière, Fabienne Proamer, Jean-Yves Rinckel, Anita Eckly, Zihan Qu, Jinmin Miao, Zhong-Yin Zhang, Yotis A. Senis, Alexandra Mazharian

**Author notes:** Corresponding author: Alexandra Mazharian, C2VN (AMU - INSERM 1263 – INRAE 1260), Aix-Marseille Université, Campus Santé Timone, Faculté des Sciences Médicales et Paramédicales, Marseille, France.

## Abstract

The Src homology 2 (SH2) domain-containing non-transmembrane protein-tyrosine phosphatases 1 and 2 (Shp1 and Shp2) have been implicated in regulating signaling from a variety of receptors and cell types, including the thrombopoietin (Tpo) receptor Mpl in megakaryocytes (MKs) and platelets. We previously showed that deletion of Shp1 and Shp2 in the MK/platelet lineage in mice using the *Pf4-Cre* transgene/*loxP* system impairs megakaryopoiesis and thrombopoiesis. However, we also observed unexpected phenotypes including a *motheaten-like* phenotype in Shp1-deficient mice and severe myelofibrosis in mice lacking both phosphatases. To determine whether these were lineage-specific effects, we utilized the *Gp1ba-Cre* transgenic mouse to delete *loxP*-flanked Shp1 and Shp2 in mice. Bone marrow-derived MKs from these mice expressed approximately 20-25% of Shp1 and Shp2, whereas platelets contain 5-10% of each phosphatase compared with controls. Minor MK/platelet defects were observed in mice lacking either Shp1 or Shp2 alone, however mice lacking both Shp1 and Shp2 exhibited macrothrombocytopenia, mild bleeding following tail injury, and impaired GPVI-mediated platelet aggregation and Syk phosphorylation, associated with reduction GPVI and integrin α2 subunit expression. Reduced Shp1 and Shp2 expression resulting in a significant reduction in ploidy, a block in MK maturation and proplatelet-producing MKs. Tpo-mediated Ras/MAPK signaling was reduced in Shp1/2-deficient MKs. Treatment of MKs with structurally distinct Shp2 allosteric inhibitors recapitulated key aspects of the Shp2-deficient phenotype, including aberrant megakaryopoiesis and reduced Mpl signaling. Our study highlights the synergistic functions of Shp1 and Shp2 in the MK/platelet lineage, and identifies Shp2 as a potential therapeutic target in myeloproliferative neoplasms.

**Key Points:**

- Deletion of Shp1 and Shp2 in the MK/platelet lineage in mice results in macrothrombocytopenia and minor effects on platelet function.
- Defects can be partially explained by reduced Mpl signaling and aberrant megakaryopoiesis in the absence of Shp2 activity.

## Introduction

Platelets, or thrombocytes are small, anucleate blood cells derived from megakaryocytes (MKs) that are essential for hemostasis and thrombosis.^1^ MK development, or megakaryopoiesis and platelet production, or thrombopoiesis are tightly regulated processes that take place in the bone marrow (BM), spleen and lungs, but the molecular mechanisms regulating these inter-related processes remain incompletely defined.^2,3^ The primary driver of megakaryopoiesis, which encompasses differentiation of hematopoietic stem and progenitor cells (HSPCs) to MKs, proliferation, endomitosis and maturation is the cytokine thrombopoietin (Tpo) produced by the kidneys, liver and BM acting on its receptor, myeloproliferative leukemia protein (Mpl) expressed throughout the MK/platelet lineage.^4^ Tpo does not however explain which MKs undergo proplatelet formation and thrombopoiesis, which occurs when MKs come in contact with sinusoidal blood vessels. Several other cytokines and growth factors, extracellular matrix (ECM) proteins and chemokines are also required for optimal megakaryopoiesis and thrombopoiesis, as evidenced in mice lacking either Tpo or Mpl, which still produce MKs and platelets, albeit at much lower levels than normal mice.^5,6^

Tpi signaling through its receptor Mpl is a central regulator of megakaryopoiesis and platelet production. Upon ligand binding, Mpl undergoes conformational changes that activate three primary signaling pathways : Janus kinase 2 (JAK2)/Signal transducer and activator of transcription (STAT), Ras/mitogen-activated protein kinase (MAPK) and phosphatidylinositol 3-kinase (PI3K)/AKT, all of which culminate in nuclear translocation of signaling/transcription factors and gene regulation.^7^ These pathways act in a coordinated manner to control both the expansion and functional maturation of megakaryocytes, thereby linking extracellular Tpo stimulation to the cellular processes governing thrombopoiesis. Cytoskeletal remodeling via the RhoA/Rho-associated protein kinases (ROCK) pathway is also an integral part of megakaryopoiesis and thrombopoiesis, notably during endomitosis^8^, but the molecular link with the Mpl receptor is undefined. Clonal gain-of-function mutations in the Mpl/JAK2/STAT pathway in HSPCs are the most common causes of myeloproliferative neoplasms (MPN)^9^, characterized by increase MK/platelet production (essential thrombocythemia, ET), increased red blood cell production (polycythemia vera, PV) and associated fibrotic deposits (primary myelofibrosis, PMF).^10–12^ Notably, the valine 617 phenylalanine (V617F) mutation in the pseudo-kinase domain of JAK2, resulting in increased catalytic active is the most prevalent, found in 60% of MPN cases, followed by C-terminal deletions in calreticulin, resulting in the release of truncated calreticulin and it acting as a ligand of Mpl in 30% of the cases, and mutations in Mpl resulting in increased expression in <5% of the cases.^13^ JAK2 and MAPK kinase (MEK) inhibitors are effective in some instances of MPN^14^, however there is a tendency of resistance and relapse, requiring alternative and improved therapeutics.

The Src homology 2 (SH2) domain-containing non-transmembrane protein-tyrosine phosphatases (PTPs) 1 and 2 (Shp1 and Shp2) are involved in hematopoietic cell differentiation, proliferation, function and survival, including megakaryopoiesis and thrombopoiesis.^15^ Intra-and inter-molecular interactions of the tandem SH2 domains with the catalytic domain and phospho-tyrosine binding partners regulate the activity and compartmentalization of both phosphatases, respectively. Tandem phospho-tyrosine residues in the C-terminal tails of Shp1 and Shp2 are implicated in substrate binding and regulation.^16^ Despite high structural similarities, the two phosphatases have distinct biological functions. Shp1, encoded by the *PTPN6* gene, is primarily expressed in hematopoietic cells and is a negative regulator of immune receptor signaling, particularly the immunoreceptor tyrosine-based activation motif (ITAM)/Spleen tyrosine kinase (Syk)/phospholipase pathway.^17^ Shp2, encoded by the *PTPN11* gene, is ubiquitously expressed and is a positive regulator of cytokine and growth factor signaling, specifically the Ras/MAPK and PI3K/AKT pathways.^18^ However, their functions go beyond these pathways and are context dependent. Downstream substrates and effectors of both phosphatases remain elusive.

We previously showed that targeted deletion of Shp1 and Shp2 in the MK/platelet lineage using the *Pf4-Cre* transgene/*loxP* system impairs megakaryopoiesis and thrombopoiesis in mice.^15^ However, we also found that these mice exhibited unexpected, non-MK/platelet-related defects, including a *motheaten-like* phenotype in Shp1-deficient mice, and severe myelofibrosis in mice lacking both phosphatases. Intriguingly, the *motheaten-like* phenotype was lost in Shp1/2 double-deficient mice, suggesting that Shp1 and Shp2 have opposing effect in this pathology. To determine whether these were in fact MK/platelet lineage-specific effects, we utilized the *Gp1ba-Cre* transgenic mouse to delete *loxP*-flanked Shp1 and Shp2 in mice. In contrast to *Pf4-Cre* transgenic mice, conditional deletion of Shp1 or Shp2 using the *Gp1ba-Cre* transgene had little if any detectable effect on megakaryopoiesis or thrombopoiesis in these mice under steady state conditions. However, deletion of both Shp1 and Shp2 using *Gp1ba-Cre* resulted in macrothrombocytopenia and aberrant MK development and maturation, due to a block in Mpl-mediated Ras/MAPK signaling. GPVI-induced platelet aggregation was also impaired due to reduced GPVI and α2 integrin subunit expression. Similar effects were seen with the structurally-distinct allosteric Shp2 inhibitors, SHP099 and RMC-4550.^19^ Findings demonstrate synergistic effects of deleting Shp1 and Shp2 in the MK/platelet lineage in mice by disrupting distinct signaling pathways, and highlight Shp2 as a potential therapeutic target in MPN.^20,21^

## Materials and Methods

### Mouse models

All mice used were on a C57BL/6 background. *Ptpn6*^fl/fl,^ *Ptpn11*^fl/fl^ and *Gp1ba-Cre^+/KI^* mice were generated, as previously described.^22–24^ MK/Platelet specific Shp1 and Shp2 knockout (KO) mice were generated by crossing *Ptpn6*^fl/fl^ and *Ptpn11*^fl/fl^ mice with the *Gp1ba-Cre* transgenic deleter mouse *Gp1ba-Cre^KI/+^* mice. *Gp1ba-Cre ^KI/+^* were used as control mice (CT) **(Supplemental Figure S1)**. All animal experiments were conducted in accordance with the CREMEAS Committee on the Ethics of Animal Experiments of the University of Strasbourg (Permit Number: E67-482-10, Project approval number: APAFIS#28221-2020111714459066).

### Inhibitors

SHP099 and RMC-4550 Shp2 allosteric inhibitors were purchased from MedChemExpress. M029 is a covalent allosteric Shp1 inhibitor (Z.Q. and Z.Y.Z, manuscript in preparation) and F2Ac is a reversible active site directed Shp1 inhibitor (J.M. and Z.Y.Z., manuscript in preparation.

### Platelet preparation and functional assays

Blood was collected from the aorta of anesthetized mice into 1/10 acid-citrate-dextrose (ACD) anticoagulant. Platelet counts were measured from peripheral blood samples using an automated hematology analyzer (Element HT5 counter, Heska company). Washed platelets were prepared as previously described.^25^ Platelet counts were normalized and used for aggregation (2 x 10^8^/ml) or biochemical analysis (5 x 10^8^/ml). Platelet aggregation was measured using the lumi-aggregometer APACT®4004. Surface glycoprotein expression was measured in whole blood by flow cytometry using fluorescein isothiocyanate (FITC)- conjugated antibodies. Resting and activated platelets were fixed and stained with anti-P-selectin antibody.

### Megakaryocyte culture and functional assays

Mature MKs from mouse BM were cultured and analysed, as previously described.^26^ BM cells were isolated from femora, tibiae and iliac crests of C57BL6 mice for pharmacological study and *Gp1ba-Cre* mice for other experiments by flushing the bones with Dulbecco’s Modified Eagle Medium (DMEM) and successively passed through 21-, 23-, and 25-gauge needles to obtain a single-cell suspension. After centrifugation, BM cells were subjected to immunomagnetic lineage-negative (Lin−) selection (StemCell Technologies) and were cultured for 3 days at 37°C 5% CO_2_ at a concentration of 1x10^6^ cells/well in DMEM supplemented with 10% Fetal Bovine Serum (FBS), 50 ng/ml Tpo, 100 U/ml Hirudin, 1% PSG.^26^ MK ploidy was assessed after 3 days of culture, using FITC-coupled anti-CD41 antibody and DNA staining with propidium iodide (PI). Samples were acquired using the BD LSRFortessa X-20 flow cytometer and analyzed using BD FACSDiva software. The ability of MKs to form proplatelets was assessed *in vitro*, and *ex vivo* using the explant assay, as previously described.^27^

### Immunoblotting

Platelet and MK whole cell lysates (WCLs) were prepared and analyzed by automated capillary-based immunoassay (ProteinSimple Jess), prepared according to manufacturer’s instructions, as previously described.^28^ The High Dynamic Range profile was used for chemiluminescent and fluorescent multiplexing signal detection. Optimized antibody dilutions and sample concentrations used are provided in **Supplemental Table 1**.

### Immunohistochemistry

Spleens from KO and litter-matched CT mice were fixed in buffered formalin and embedded in paraffin. Sections (5 μm) were H&E stained and examined by light microscopy using a 40× objective.

### Electron microscopy

Maturation stage of MKs was assessed by electron microscopy (EM). BM samples obtained by flushing mouse femora with 0.1 M sodium cacodylate buffer were fixed in 2.5% glutaraldehyde and embedded in Epon as previously described.^29^ Thin sections were stained with uranyl acetate and lead citrate, and examined under a JEOL 2100Plus transmission electron microscope at 120 kV (Jeol, Tokyo, Japan). MKs at stages I, II and III were identified using distinct ultrastructural characteristics: stage I, a cell 10–15 μm in diameter with a large nucleus; stage II, a cell 15–30 μm in diameter containing platelet-specific granules; stage III, 40 µm, a MK containing a well-developed demarcation membrane system defining cytoplasmic territories and a peripheral zone. Samples from at least three mice of each genotype were examined in each case.

### Platelet recovery assay

An intraperitoneal injection of anti-GPIbα antibody (Emfret) at 2 mg/kg is performed at D0. Blood samples were collected and platelet counts measured prior to injection, and at D5, D10, D15 and D20.^30^ Platelet recovery kinetics were analyzed using two-way ANOVA with appropriate post hoc tests.

### HSPC colony-forming unit (CFU) assay

Cells are obtained by flushing and were resuspended in IMDM and plated at a density of 1×10⁴ cells per dish in methylcellulose-based medium (MethoCult™ GF M3434, Stemcell Technologies) containing appropriate cytokines to support multilineage hematopoietic colony formation. Cultures were incubated at 37 °C with 5% CO₂ in a humidified incubator for 10–14 days. Colonies were scored under an inverted microscope and classified based on morphological criteria into CFU-G (granulocyte), CFU-M (macrophage), CFU-GM (granulocyte-macrophage), BFU-E (erythroid), and CFU-GEMM (granulocyte, erythrocyte, macrophage, megakaryocyte). Only colonies containing ≥50 cells were counted. Results were expressed as the mean colony number ± SEM from n = 3 independent experiments.

### Statistical analysis

Statistical analysis was performed using GraphPad Prism version 9 software. Data were expressed as mean ± standard error mean (SEM). Statistical significance was analyzed by one-or two-way ANOVA followed by the appropriate *post hoc* test, or Kruskal-Wallis test for nonparametric data, a Bonferroni correction was applied for multiple comparisons, as indicated in figure legends. P-values < 0.05 were considered statistically significant.

## Results

### Deletion of Shp1 and Shp2 in the MK/platelet lineage

To validate the efficiency of protein ablation of Shp1 and Shp2 proteins in MKs/platelets using the *Gp1ba-Cre* transgenic mouse strain **(Supplemental Figure S1)**, we employed a quantitative capillary-based immunoassay. This was performed on cells derived from *Ptpn6^fl/fl^*;*Gp1ba-Cre^+/KI^* (Shp1 KO), *Ptpn11^fl/fl^*;*Gp1ba-Cre^+/KI^* (Shp2 KO) and *Ptpn6^fl/fl^*;*Ptpn11^fl/fl^*;*Gp1ba-Cre^+/KI^* double-knockout (DKO) mice, with *Gp1ba-Cre^+/KI^* littermates serving as controls.

We firstly evaluated Shp1 and Shp2 levels in MK-sorted progenitors (MKPs) in order to obtain a pure population of matures MKs. Under these conditions, we quantified protein loss of Shp1 and Shp2 in MKs, as shown in **Figure 1Ai-iii,** with significant 70% and 82% reduction of Shp1 protein in Shp1 KO and Shp1/2 DKO mice, respectively and 78% and 75% reduction of Shp2 protein in Shp2 KO and Shp1/2 DKO mice, respectively compared to controls.

**Figure 1.**
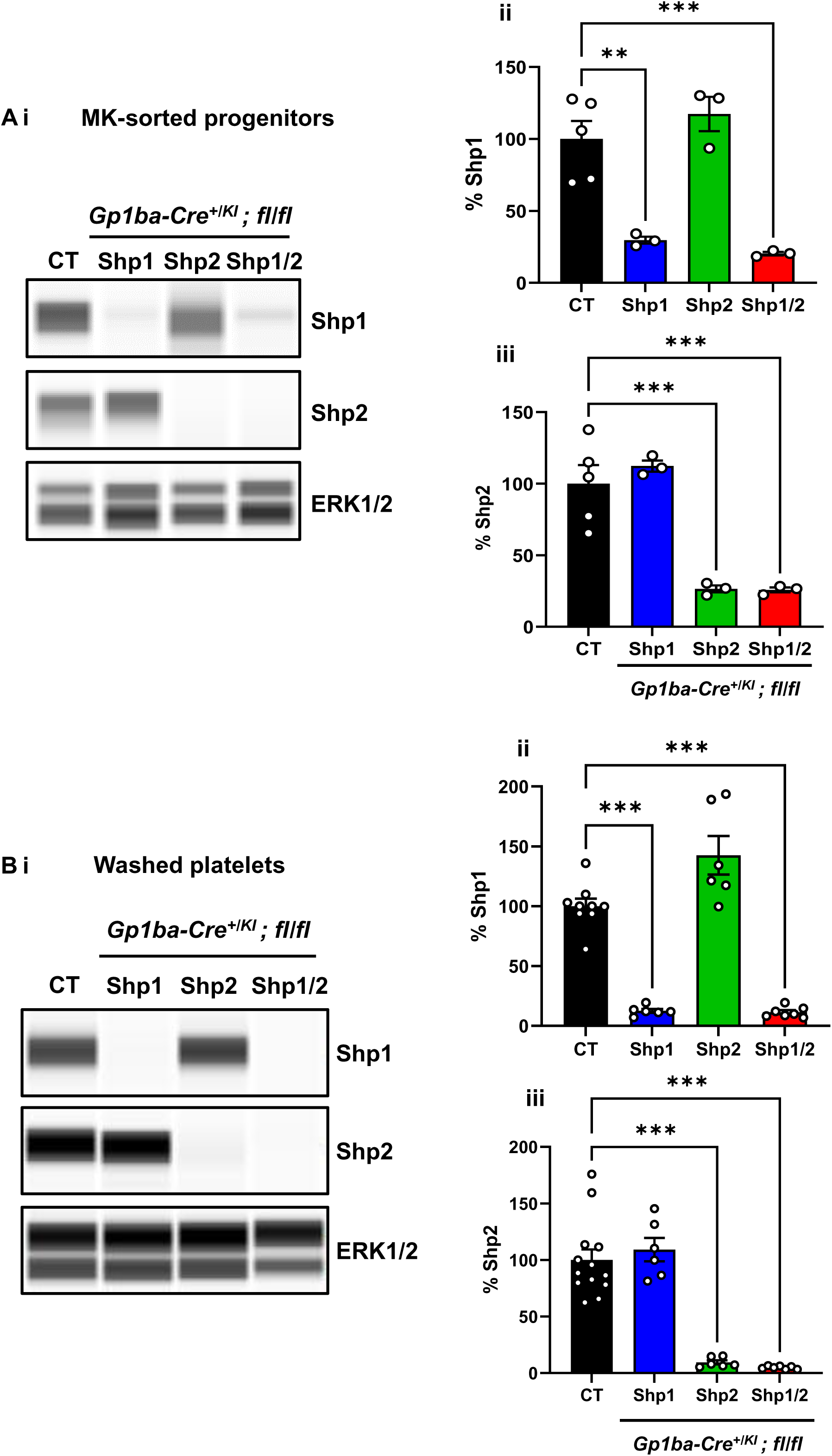
Protein expression of Shp1 and Shp2 in megakaryocytes and platelets from *Gp1ba-Cre* mice. **(i)** Representative blots of Shp1 and Shp2 expression levels from CT, Shp1, Shp2 and Shp1/2 DKO mice by capillary immunoassays with the respective antibodies in **(A)** sorted-MK progenitors and **(B)** washed platelets. Percentage of residual level of **(ii)** Shp1 and **(iii)** Shp2 in Shp1 KO (blue), Shp2 KO (green) and Shp1/2 DKO (red) mice. n = 3-6 mice per genotype. Mean ± SEM; one way-ANOVA; ** *P* < 0.01, *** *P* < 0.001.

We next evaluated protein levels in washed platelets and found efficient deletion in Shp1 KO and Shp2 KO platelets, with a significant 90% and 92% reduction of Shp1 protein in Shp1 KO and Shp1/2 DKO mice respectively, and 95% and 96% reduction of Shp2 protein in Shp2 KO and Shp1/2 DKO mice respectively, compared to controls (**Figure 1Bi-iii)**. In addition, there was no upregulation of Shp1 in Shp2-deficient MKs/platelets and vice versa. These results indicate that the *Gp1ba-Cre* transgene efficiently deleted *loxP*-flanked *Ptpn6* and *Ptpn11*, resulting in the significant loss of Shp1 and Shp2 in MKs/platelets.

### Macrothrombocytopenia and increased bleeding in Shp1/2 DKO mice

Deletion of Shp1 or Shp2 using the *Gp1ba-Cre*-mouse strain did not affect platelet counts and volumes as shown in **Figure 2Ai-ii and Supplemental Table 2**. However, Shp1/2 DKO mice were macrothrombocytopenic, with a 40% reduction in platelet count and 15% increase in platelet volume **(Figure 2Ai-ii)**. Monocyte, neutrophil and eosinophil counts were marginally increased in Shp1/2 DKO mice, suggesting possible immune responses. **(Supplemental Table 2)**.

**Figure 2.**
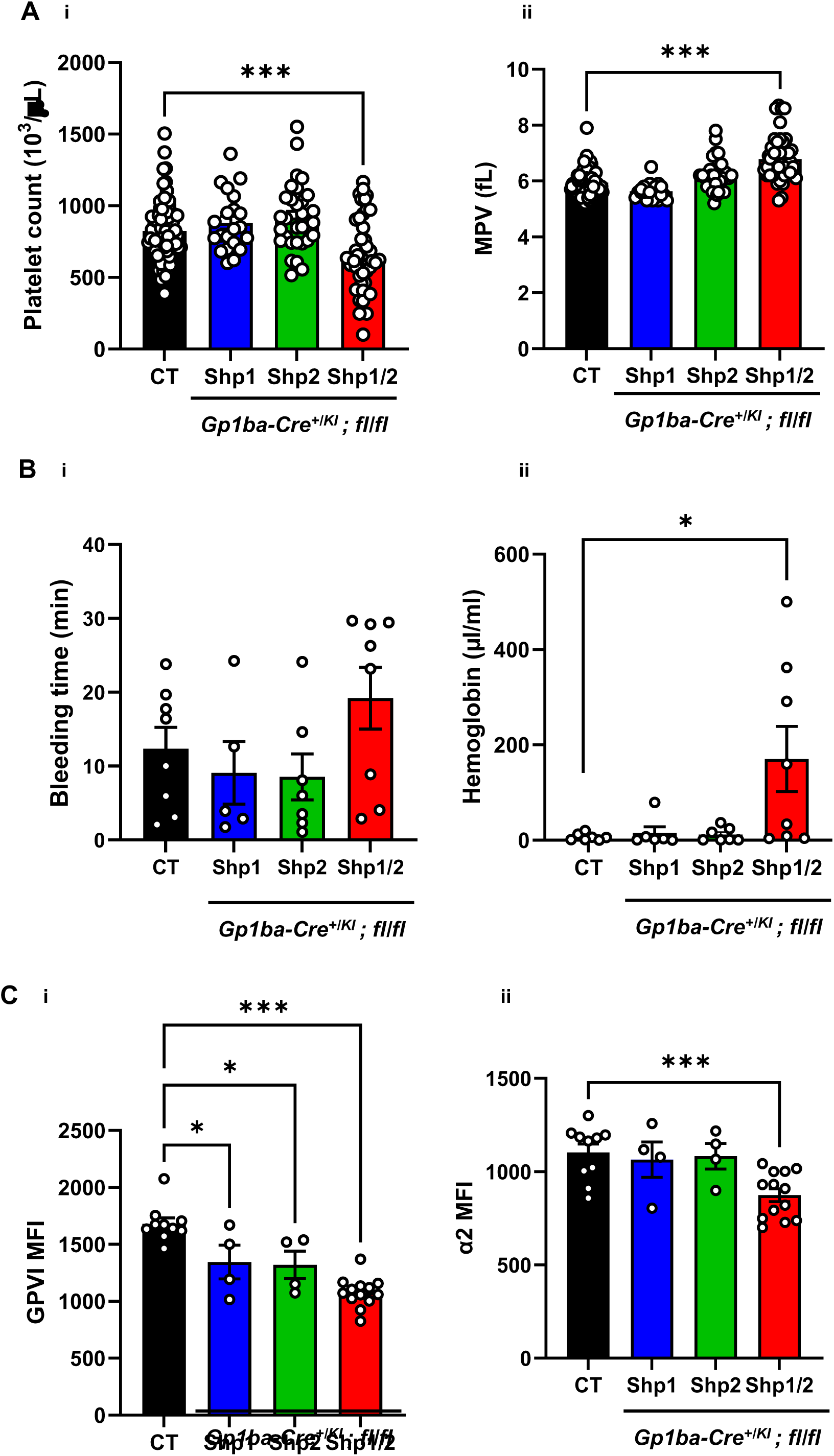
Macrothrombocytopenia and mild increased bleeding in Shp1/2 DKO mice. **(A)** Platelet counts **(i)** and platelet volumes **(ii)** of control (CT) (n=75), Shp1 KO (n=21), Shp2 KO (n=29) and Shp1/2 DKO (n=46) mice. Mean ± SEM; one way-ANOVA; *** *P* < 0.001. **(B) (i)** Cumulative bleeding time was recorded **(ii)** and the volume of blood loss was measured over 30 minutes. Mean ± SEM; one way-ANOVA; * *P* < 0.05. **(C)** Mean fluorescence intensity (MFI) of platelet **(i)** GPVI and **(ii)** α2 integrin expression in CT (n=10), Shp1 KO (n=4), Shp2 KO (n=4) and Shp1/2 DKO (n=13) mice. Mean ± SEM; one way-ANOVA; *** *P* < 0.001.

The primary function of platelets is to prevent blood loss following injury. Therefore, to assess the impact of the loss of Shp1 and Shp2 on hemostasis, we performed tail bleeding assays. Although bleeding time was not significantly different between Shp1/2 DKO and control mice (**Figure 2Bi)**, Shp1/2 DKO exhibited a significant increase in blood loss as reflected by higher hemoglobin levels (**Figure 2Bii),** indicating impaired hemostasis.

### Aberrant functional responses of Shp1/2-deficient platelets

We next investigated platelet receptor expression by flow cytometry. The collagen activation receptor GPVI receptor expression was reduced in Shp1- and Shp2-deficient platelets compared with controls. However, the collagen activation receptor GPVI and the α2 subunit of the collagen integrin α2β1 were reduced by 25% and 13%, respectively, in Shp1/2-deficient platelets **(Figure 2Ci-ii and Supplemental Table 3)**. All other major surface receptors analyzed were unaltered in these platelets **(Supplemental Figure S2 and Supplemental Table 3)**.

We next checked platelet aggregation by light transmission aggregometry. Shp1- and Shp2-deficient platelets responded normally to all agonists tested (**Figure 3A-C)**. However, consistent with reduced GPVI and α2 surface expression, Shp1/2-deficient platelets exhibited a marginal reduction in aggregation to a low and intermediate concentration of the GPVI-specific agonist CRP (1 and 3 μg/mL), and collagen (3 μg/mL), which signals via GPVI and binds with high affinity to the integrin α2β1 **(Figure 3Ai-ii)**. However, they responded normally to thrombin (0.06 and 0.1 U/ml) **(Figure 3Bi-ii),** antibody-mediated cross-linking of the hem-ITAM-containing podoplanin receptor CLEC-2 (3 μg/mL) and ADP (3 μM) **(Figure 3Ci-ii)**.

**Figure 3.**
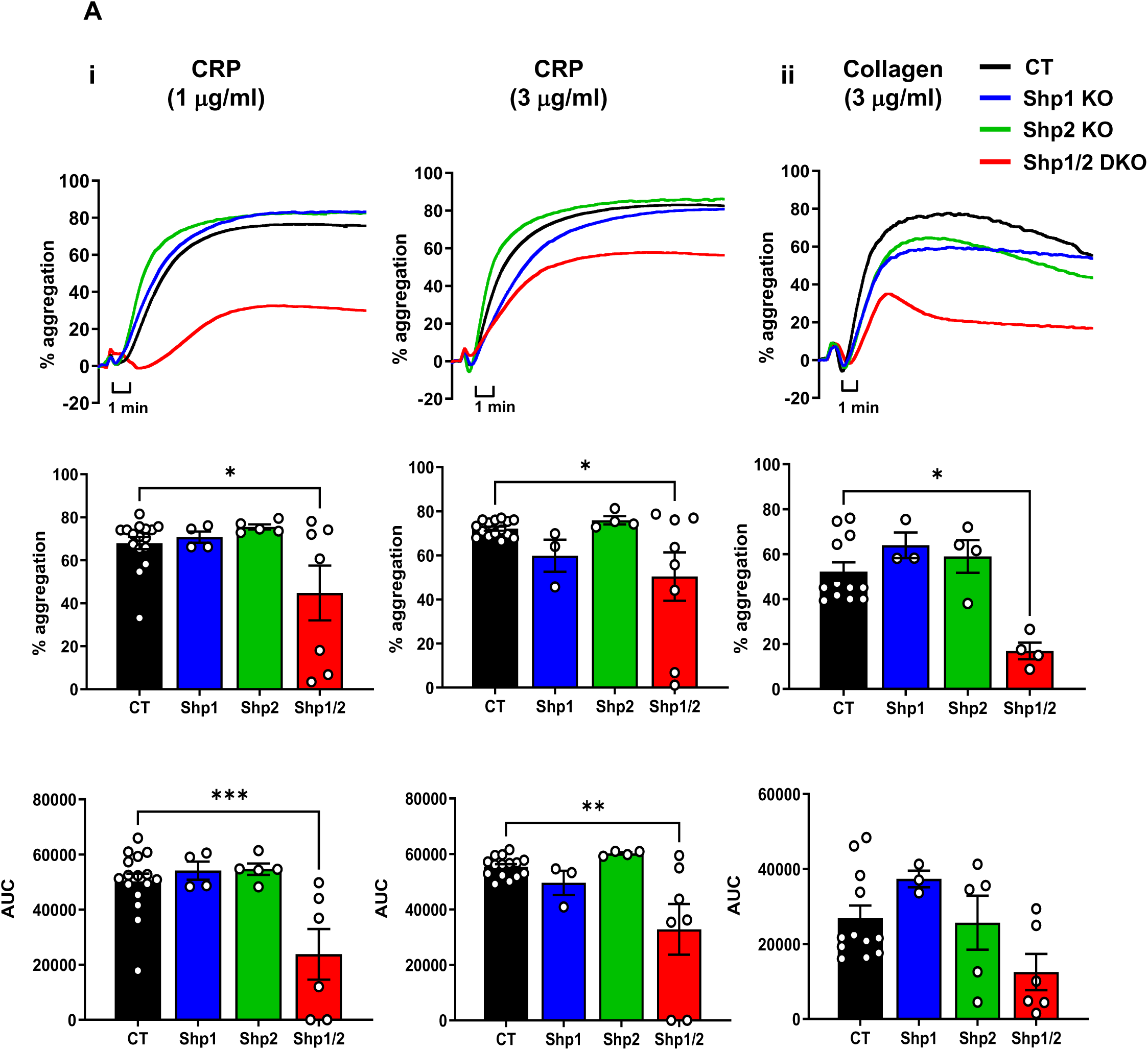
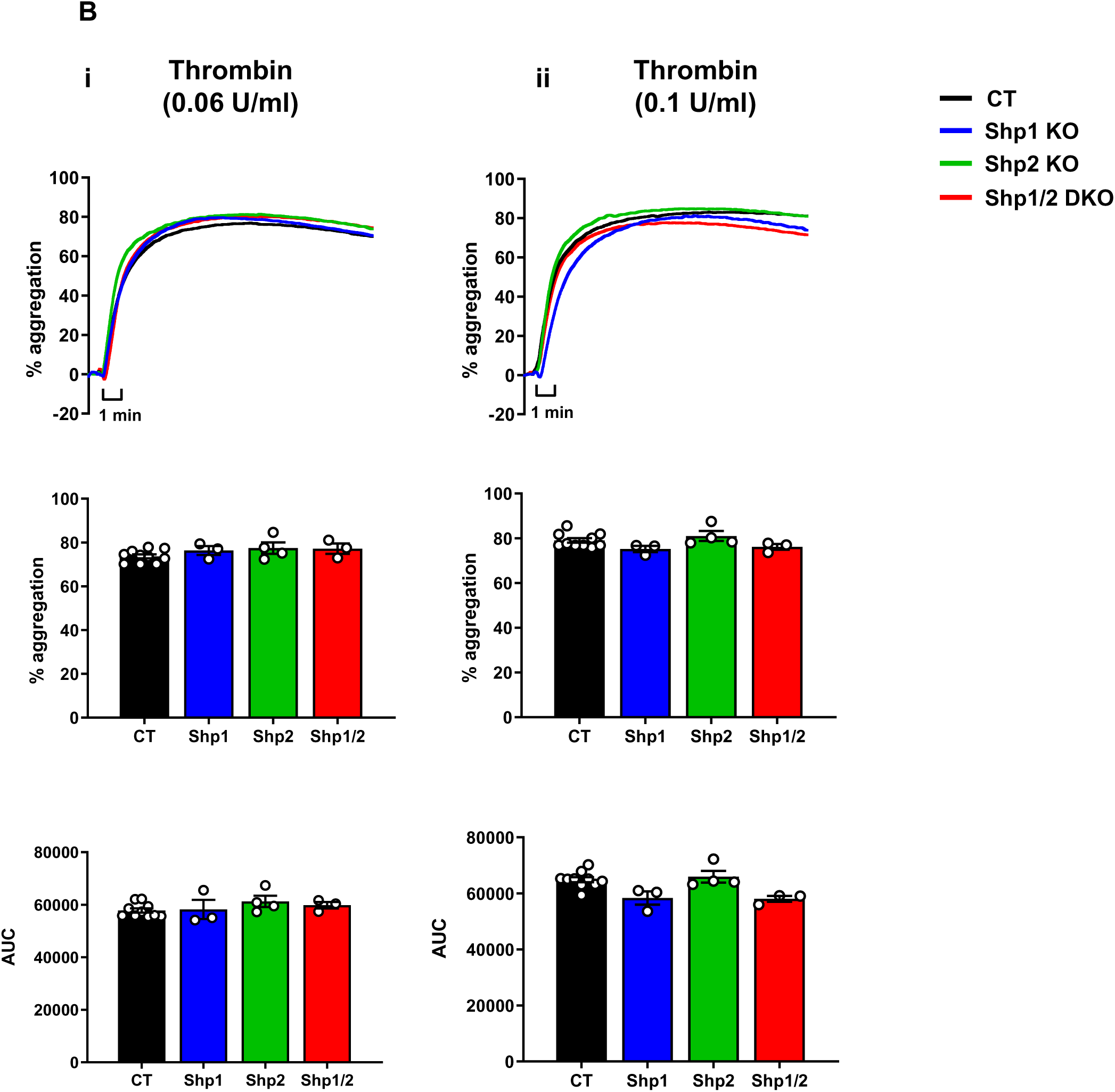
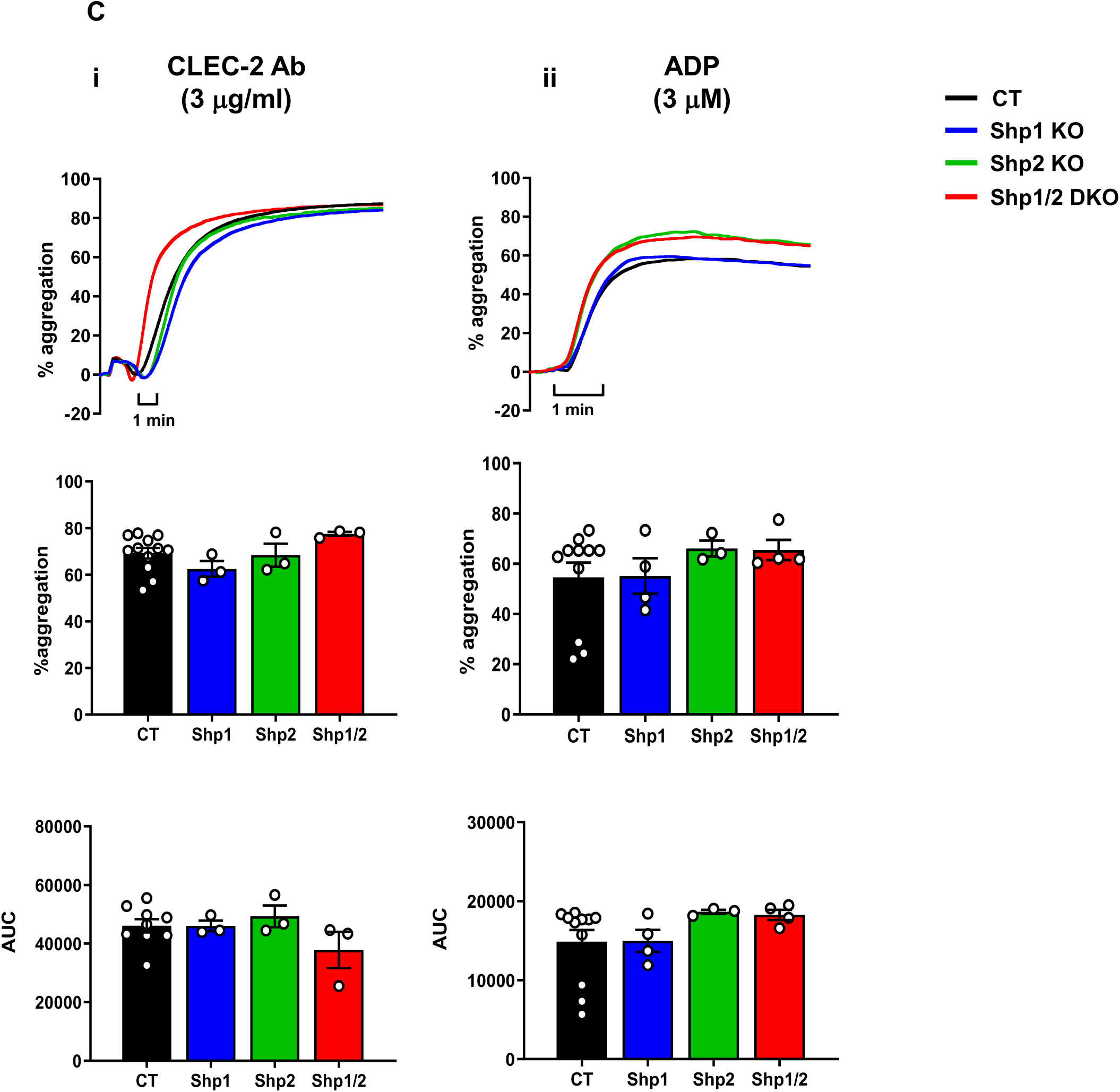
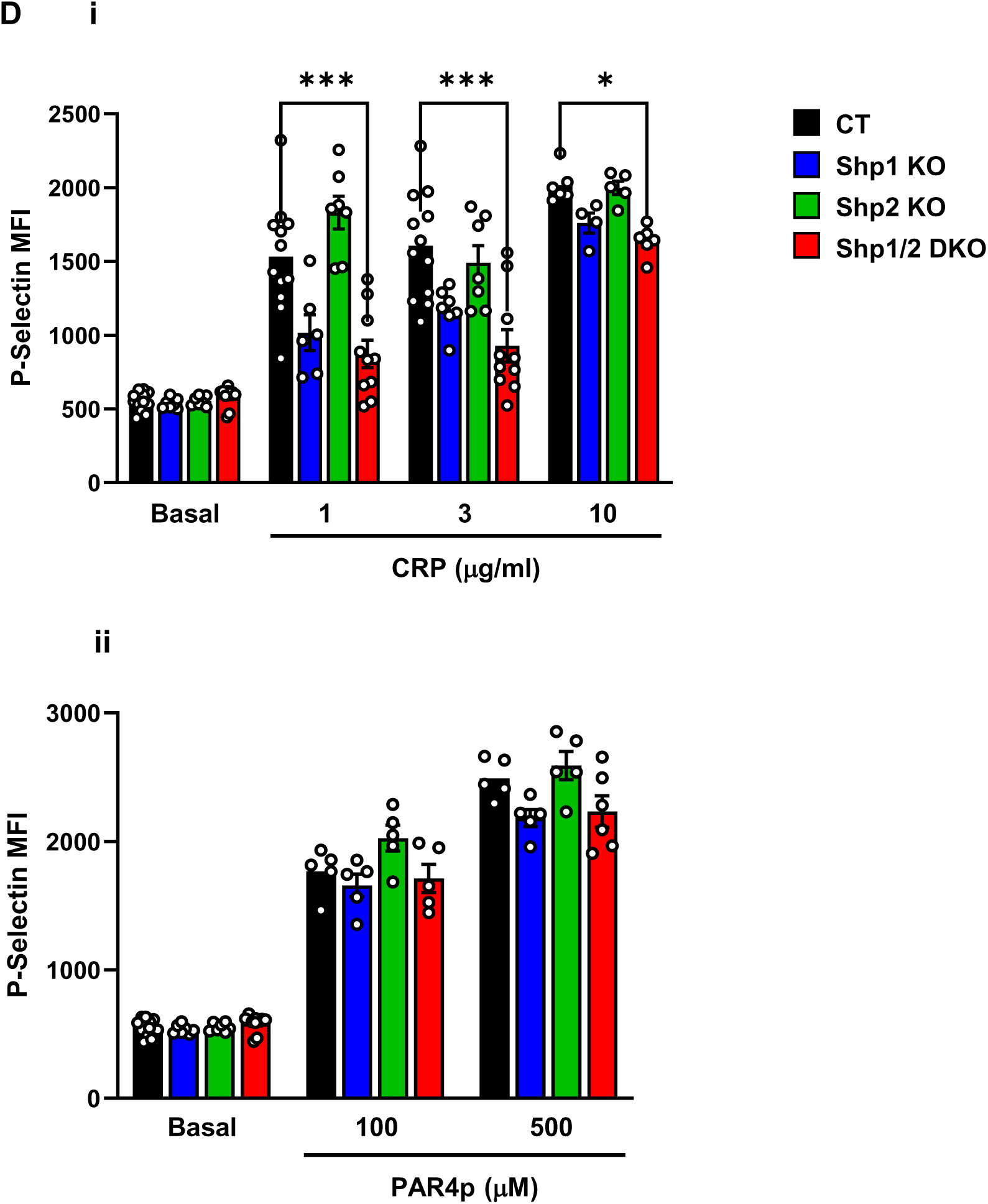
Aberrant platelet function in Shp1/2 DKO mice. **(A-C)** Aggregation of washed platelets were measured by lumi-aggregometry in response to agonists indicated. Representative traces from independent experiments included in the quantitative analysis are shown. n= 4-8 mice/genotype per condition, percentage of aggregation at 5 minutes and area under the curve (AUC) quantification. Mean ± SEM, n = 5-17 mice/genotype per condition, one way-ANOVA, * *P* < 0.05. **(D)** Mean fluorescence intensity (MFI) of P-selectin expression of control (CT), Shp1 KO, Shp2 KO and Shp1/2 DKO platelets in whole blood in response to **(i)** 1 and 3 μg/mL CRP and **(ii)** 100 and 500 μM PAR4 peptide (PAR4p). Mean ± SEM, n = 5-9 mice/genotype per condition, two way-ANOVA; * *P* < 0.05, *** *P* < 0.001.

We also measured P-selectin surface expression in whole blood as a marker of α-granule secretion following platelet activation. Consistent with reduced GPVI expression, Shp1/2-deficient platelets showed markedly reduced P-selectin exposure in response to the GPVI-specific agonist CRP (1, 3, and 10 μg/mL) **(Figure 3Di)**, whereas P-selectin expression in response to PAR-4 peptide (100 and 500 μM), which activates platelets through thrombin receptor signaling, was comparable to controls **(Figure 3Dii)**.

### Aberrant ITAM signaling in Shp1- and Shp2-deficient platelets

Since Shp1 and Shp2 have been implicated in regulating ITAM-containing receptor signaling, we investigated GPVI and CLEC-2 signaling in Shp1- and Shp2-deficient platelets. We focused on Src Family Kinases (SFK), which phosphorylate tyrosine residues within the ITAM of GPVI-associated FcR γ-chain and hemi-ITAM of the CLEC-2 receptor, and thereby enabling the recruitment and activation of Syk, which docks to both via its tandem SH2 domains, and mediates downstream effects. Activated Syk subsequently propagates the signal through phosphorylation of downstream effectors, including PLCγ2, leading to calcium mobilization and platelet activation. SFK activation was indirectly measured as *trans-*autophosphorylation of the highly conserved tyrosine residue 418 in Src (Src p-Tyr418) and Syk activation was measured as *trans*-autophosphorylation of tyrosine residues 519/520 (p-Tyr519/20), which directly correlate with activity. Western blot analysis demonstrated that Src p-Tyr418 was not altered under any conditions (**Figure 4Ai-ii).** However, Syk p-Tyr519/520 was significantly decreased in CRP–stimulated platelets from Shp1/2-deficient platelets, and marginally reduced in CLEC-2-stimulated Shp1 KO and Shp2 KO deficient platelets, (**Figure 4Bi-ii)** suggesting that Shp1 and Shp2 modulate Syk-dependent signaling downstream of GPVI and CLEC-2.

**Figure 4.**
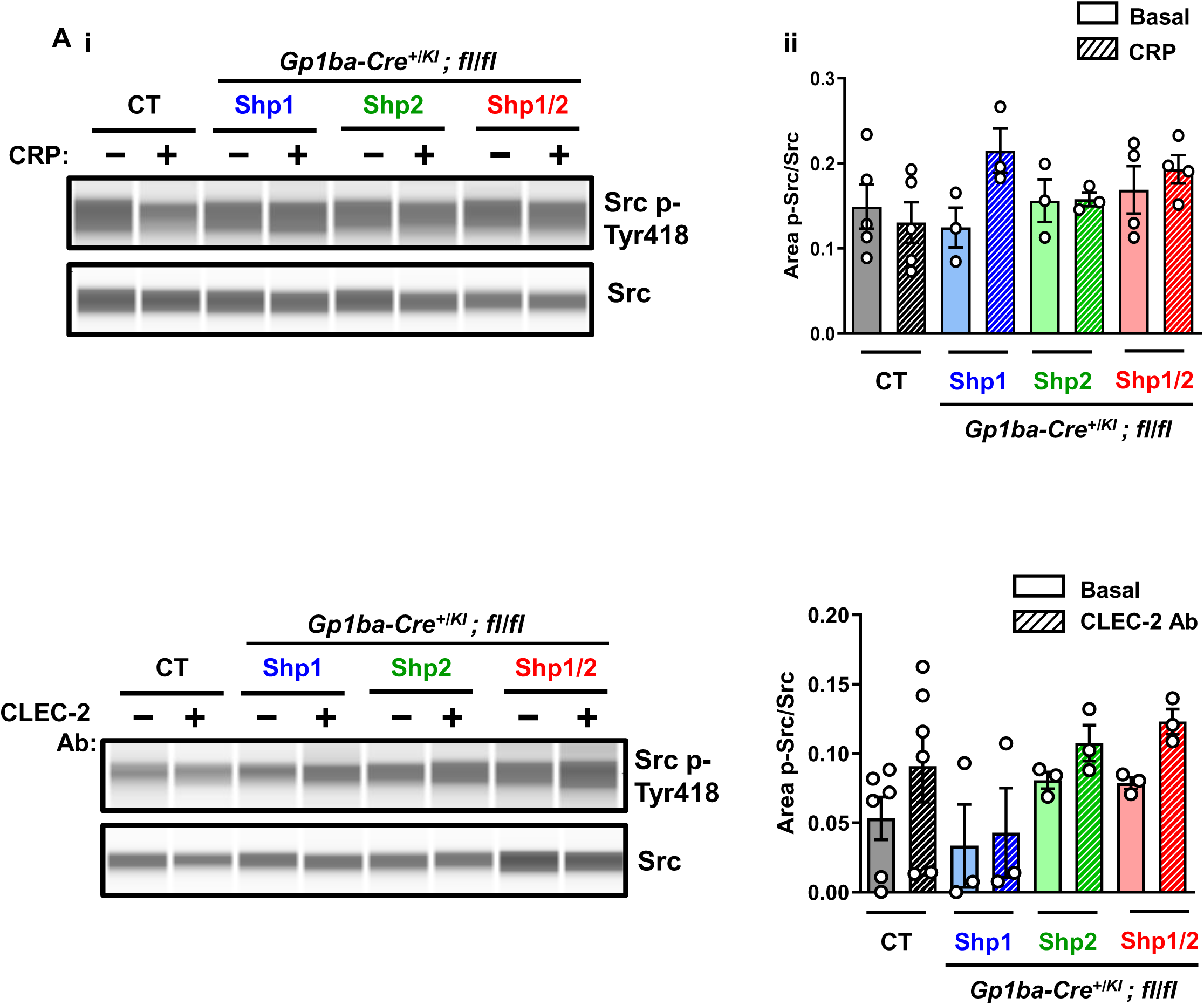
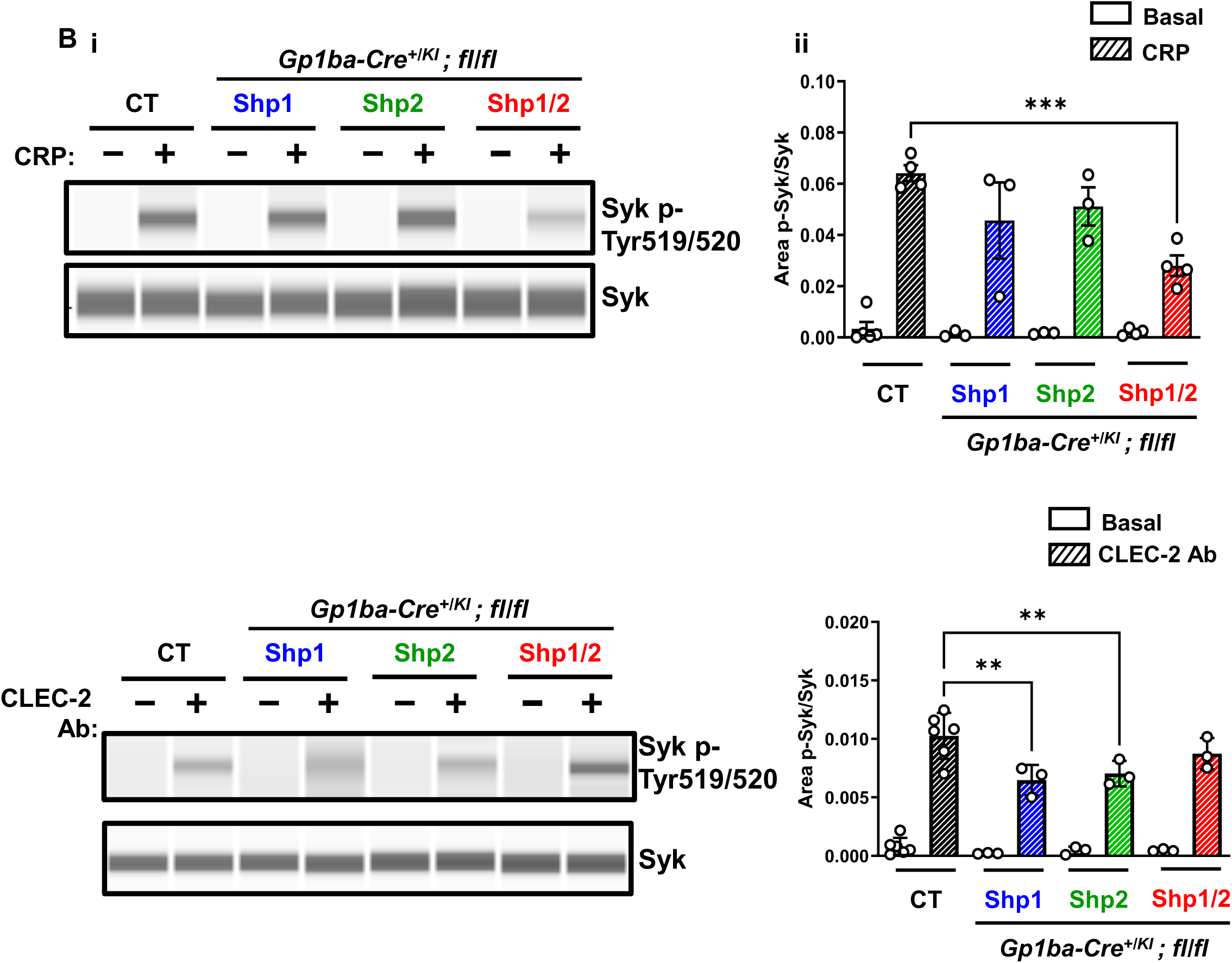
Aberrant platelet signaling in Shp1/2 DKO mice. Whole cell lysates of resting and **(A)** 10 µg/ml CRP-stimulated and **(B)** 10 µg/ml activating CLEC-2 antibody platelets from Shp1 KO, Shp2 KO and Shp1/2 DKO mice and litter-matched CT mice were western blotted with **(A)** anti-Src p-Ty418, Src total, **(B)** -Syk p-Tyr519/520 and Syk antibodies. **(i)** Representative blots of capillary-based immunoassays and **(ii)** quantification of peak areas from three independent experiments, Mean ± SEM, n = 3-4 mice/genotype; one-way ANOVA, ** *P* < 0.01, *** *P* < 0.001.

### Impaired megakaryocyte differentiation and function

To determine the cause of thrombocytopenia in Shp1/2 DKO mice, we measured platelet recovery following antibody-mediated depletion as an indicator of platelet turnover. As expected, administration of the anti-GPIbα antibody resulted in extreme and sustained thrombocytopenia, confirming efficient depletion of circulating platelets (**Supplemental Figure S3)**. No differences were observed the different phases or kinetics of platelet clearance or recovery in any of the mouse models. Interestingly, spleen size and weight were significantly increased in Shp1/2 DKO mice (**Supplemental Figure S4**), suggesting compensatory extramedullary hematopoiesis. However, HSPC colony-forming unit (CFU) assays performed on BM cells showed no significant difference in the frequency or composition of HSPCs, including MKPs between genotypes *in vitro* (**Supplemental Figure S5**).

Despite the maintenance of normal platelet turnover *in vivo*, *in vitro* analyses revealed pronounced defects in MK development, maturation and function of Shp1/2-deficient MKs (**Figure 5A-C)**. BM-derived MKs from DKO mice exhibited a severe defect in polyploidization, as shown by a significant increase in 2-8N immature MKs, and decreased 16-64N mature MKs (**Figure 5Ai-ii)**. Moreover, proplatelet formation was severely impaired, with a significantly lower percentage of MKs extending cytoplasmic projections compared to controls (**Figure 5B)**.

**Figure 5.**
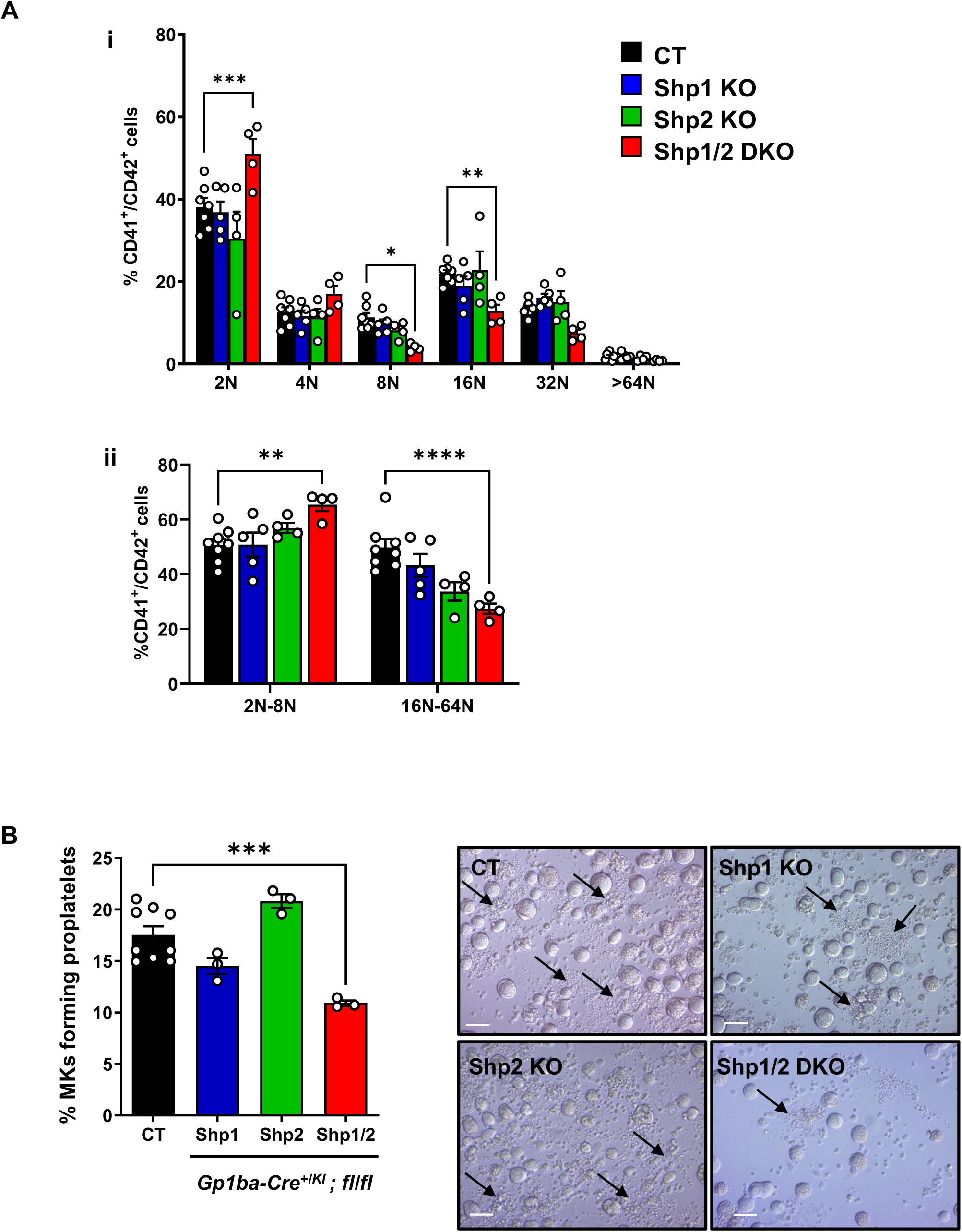
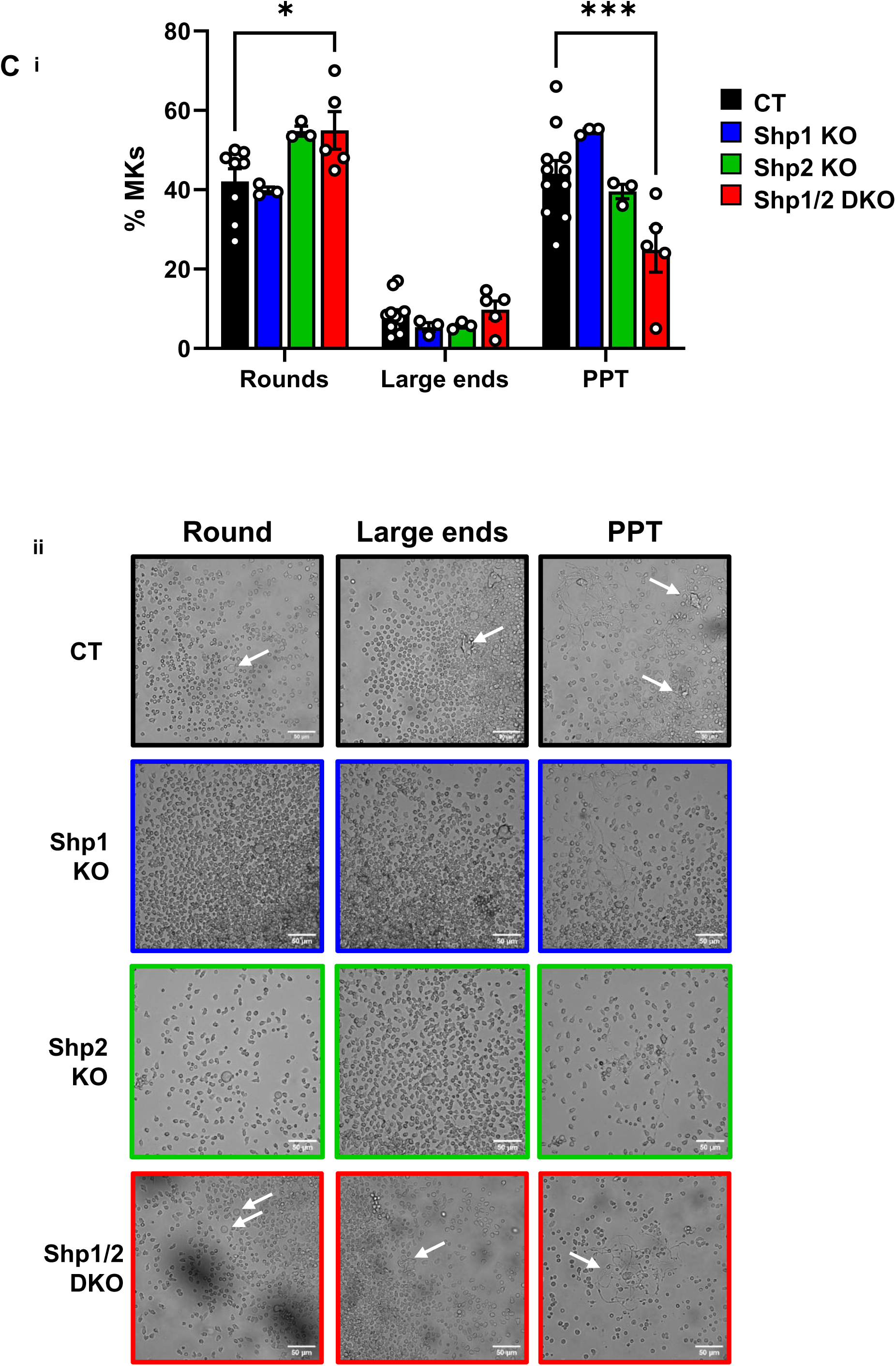
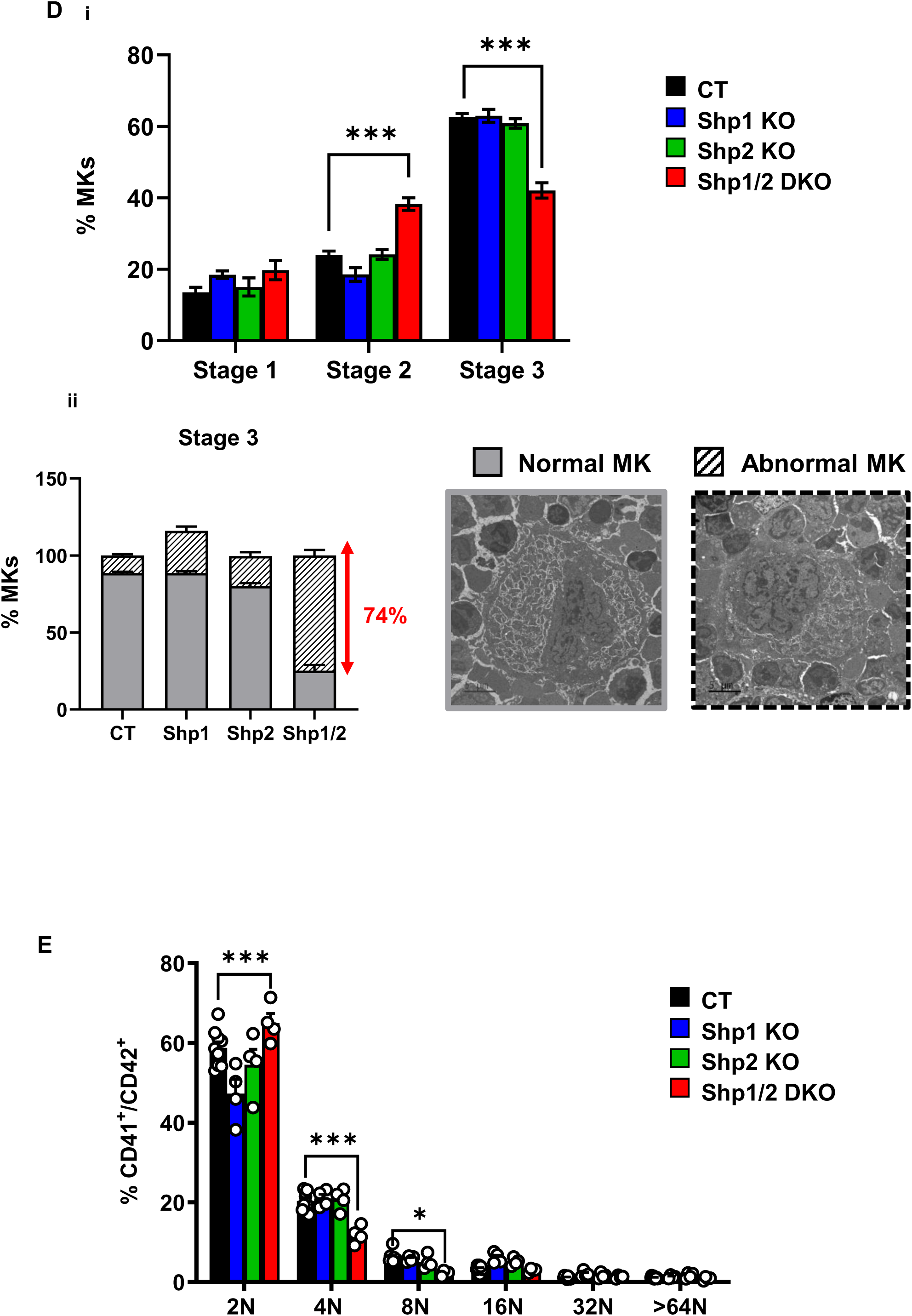
Defects in megakaryocyte maturation and *ex vivo* proplatelet formation in Shp1/2 DKO. **(A)** Mature BM-derived MKs from Shp1 KO, Shp2 KO, Shp1/2 DKO, and litter-matched CT mice were stained with propidium iodide and ploidy of cells was quantified by flow cytometry. **(i-ii)** The percentage of 2-4N and 8-128N ploidy cells was quantified (n = 4-6 mice/genotype; mean ± SEM; two-way ANOVA, * *P* < 0.05, ** *P* < 0.01, *** *P* < 0.001). **(B)** *Ex vivo* proplatelet formation. Percentage of MKs forming proplatelet was quantified in culture. Mean ± SEM, two-way ANOVA, *** *P* < 0.001. **(C)** Bone marrow explants. Proportion of MKs from Shp1 KO, Shp2 KO, Shp1/2 DKO and litter-matched CT mice extending proplatelets at 6h of observation were observed. Bars represent the mean ± SEM of six independent experiments. **(i)** Quantification and **(ii)** representatives’ images of round MKs, MKs with large ends and proplatelet (PPT) forming MKs. Scale bar: 50 µm. Mean ± SEM, one-way ANOVA, * *P* < 0.1, *** *P* < 0.001. **(D)** Classification of the MK according to their maturation stage: stage I (absence of granules), stage II (granules and developing demarcation membrane system (DMS) not yet organized), stage III (DMS organized in cytoplasmic territories). Data are reported as the percentage of the total number of MK, **(i)** of all stage MK and **(ii)** only stage 3 MK. Each bars segment represents the mean percentage ± SEM from three BM samples, *** *P* < 0.001. **(E)** Ploidy of *in situ* BM MKs measured by flow cytometry. Mean ± SEM, two-way ANOVA, * *P* < 0.1, *** *P* < 0.001.

Collectively, these data suggest that while homeostatic and compensatory thrombopoiesis is preserved *in vivo*, potentially supported by extramedullary hematopoiesis, Shp1 and Shp2 are required for efficient megakaryopoiesis *in vitro*, highlighting a critical role in late-stage megakaryopoiesis under defined conditions.

### Defective proplatelet formation and MK maturation in Shp1/2-deficient mice

To further characterize the functional consequences of Shp1 and Shp2 deficiency on megakaryopoiesis, we evaluated proplatelet formation *ex vivo* and examined MK morphology *in situ*. We first performed *ex vivo* proplatelet formation assays using BM explants. The results revealed a marked reduction in proplatelet formation in Shp1/2 DKO mice compared with controls, with an increased number of round MKs and a decreased number of MK-forming proplatelets in Shp1/2 DKO mice **(Figure 5Ci-ii)**. To assess MK maturation *in situ*, we performed EM and ploidy analysis in BM. In control mice, MK displayed characteristic features of terminal maturation, including large polyploid nuclei, demarcation membrane system (DMS) organization, and extensive cytoplasmic development. Shp1/2 DKO MK exhibited a maturation arrest, with increased frequency of stage III MK, defined by incomplete DMS formation, and limited cytoplasmic granularity **(Figure 5Di-ii).** This was further supported by flow cytometric analysis of DNA content, showing lower ploidy level (4N–8N) in Shp1/2 DKO MK (**Figure 5E)**. These findings provide further evidence of the involvement of Shp1 and Shp2 in late-stage MK maturation and proplatelet formation, absence of which results in the accumulation of immature, stage III MK in the BM and a functional failure to complete thrombopoiesis.

### Reduced Tpo signaling in Shp1/2-deficient MKs

To understand the molecular basis of impaired megakaryopoiesis in Shp1/2 DKO mice, we assessed Tpo-mediated signaling in BM-derived MKs. We previously demonstrated that Shp2 is a positive regulator of ERK1/2 downstream of Mpl. Tpo-mediated ERK1/2 activation was normal in Shp1-deficient MK and significantly reduced in Shp2- and Shp1/2-deficient MKs. However, other downstream effectors, including AKT and STAT3, showed modest or variable changes (**Figure 6A-C)**. These findings provide a mechanistic explanation for the impaired MK development and maturation observed *in vitro*, and defective Tpo signaling through the ERK1/2 pathway in Shp1/2-deficient MK, underscoring the importance of these phosphatases in late-stage megakaryopoiesis.

**Figure 6.**
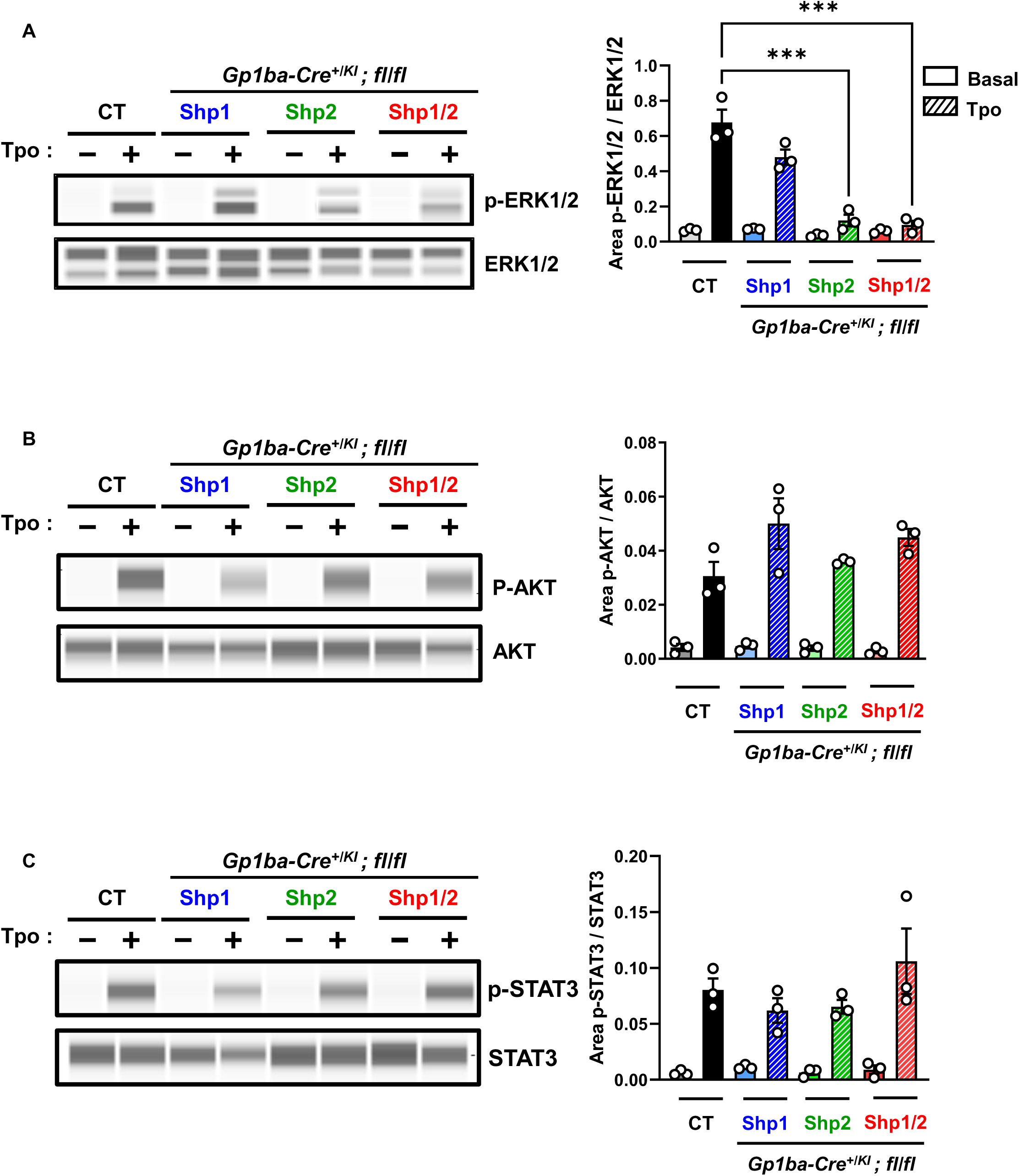
Defects in TPO signaling in Shp1/2 DKO mice. Mature BM-derived MKs from Shp1 KO, Shp2 KO, Shp1/2 DKO and litter-matched CT mice were stimulated with 50 ng/mL thrombopoietin (Tpo) for 10 min at 37°C. Whole cell lysates were western blotted with indicated antibodies **A-C**. Representative blots capillary-based immunoassays and quantification of peak areas from n = 3 independent experiments/genotype; two-way ANOVA, *** *P* < 0.001.

### Shp2 allosteric inhibitors impair megakaryopoiesis and Tpo signaling

To further delineate the respective roles of Shp1 and Shp2 in MK and Tpo signaling, we employed two pharmacologically distinct inhibitors for each phosphatase to ensure specificity and validate findings from genetically modified mouse models.

For Shp1, we used M029, a covalent allosteric inhibitor that induces conformational changes that block Shp1 activity^34^ and F2Ac, a reversible inhibitor, which directly target the catalytic site (J.M. and Z.Y.Z., manuscript in preparation). Treatment of murine BM-derived MK with these Shp1 inhibitors showed no significant effects on cell viability, proliferation and MK ploidy **(Figure 7Ai-iii)**, nor on Tpo-mediated signaling **(Figure 7B)**, correlating with what was observed in Shp1-deficient MKs. Similarly, Shp2 was inhibited using SHP099 and RMC-4550, both selective allosteric inhibitors that stabilize Shp2 in its inactive conformation^31^. Addition of Shp2 inhibitors resulted in a marginal decrease in cell viability and a marked reduction of cell proliferation, both established functions of Shp2 in other lineages **(Figure 7Ci-ii)**. Moreover, treatment with either SHP099 or RMC-4550 resulted in a significant inhibition of ploidy with a reduction in the percentage of cells achieving >8N **(Figure 7Ciii),** and a significant reduction in proplatelet formation **(Figure 7Civ)**. This was accompanied by a severe impairment in Tpo-mediated ERK1/2 and AKT phosphorylation **(Figure 7D)**. This dual-inhibitor profiling reveals that Shp2, but not Shp1, is critical for Tpo-driven MK proliferation, survival, endomitosis and Mpl signaling, highlighting Shp2 as a key regulator of late stage megakaryopoiesis and suggest that its inhibition disrupts thrombopoietic pathways.

**Figure 7.**
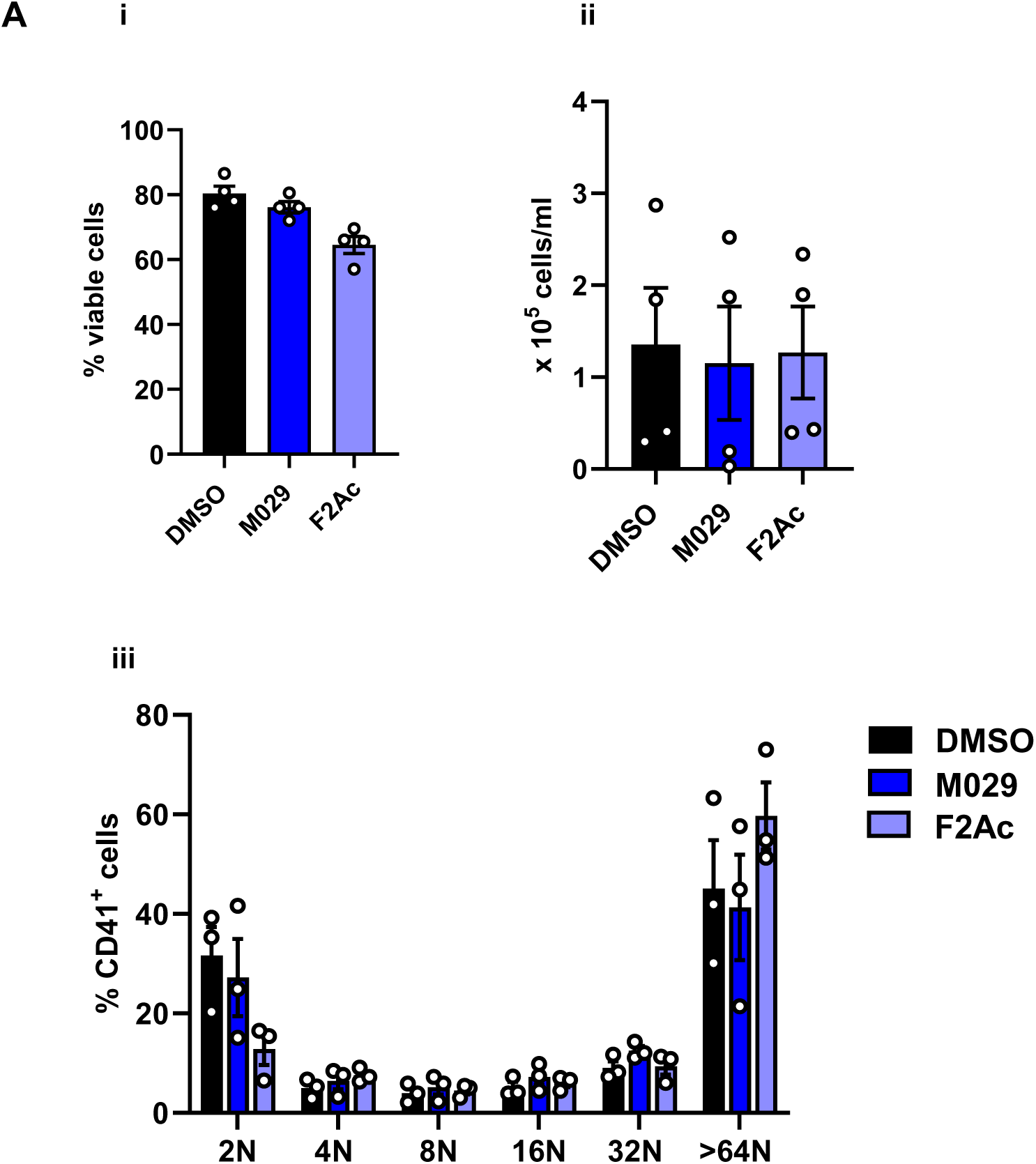
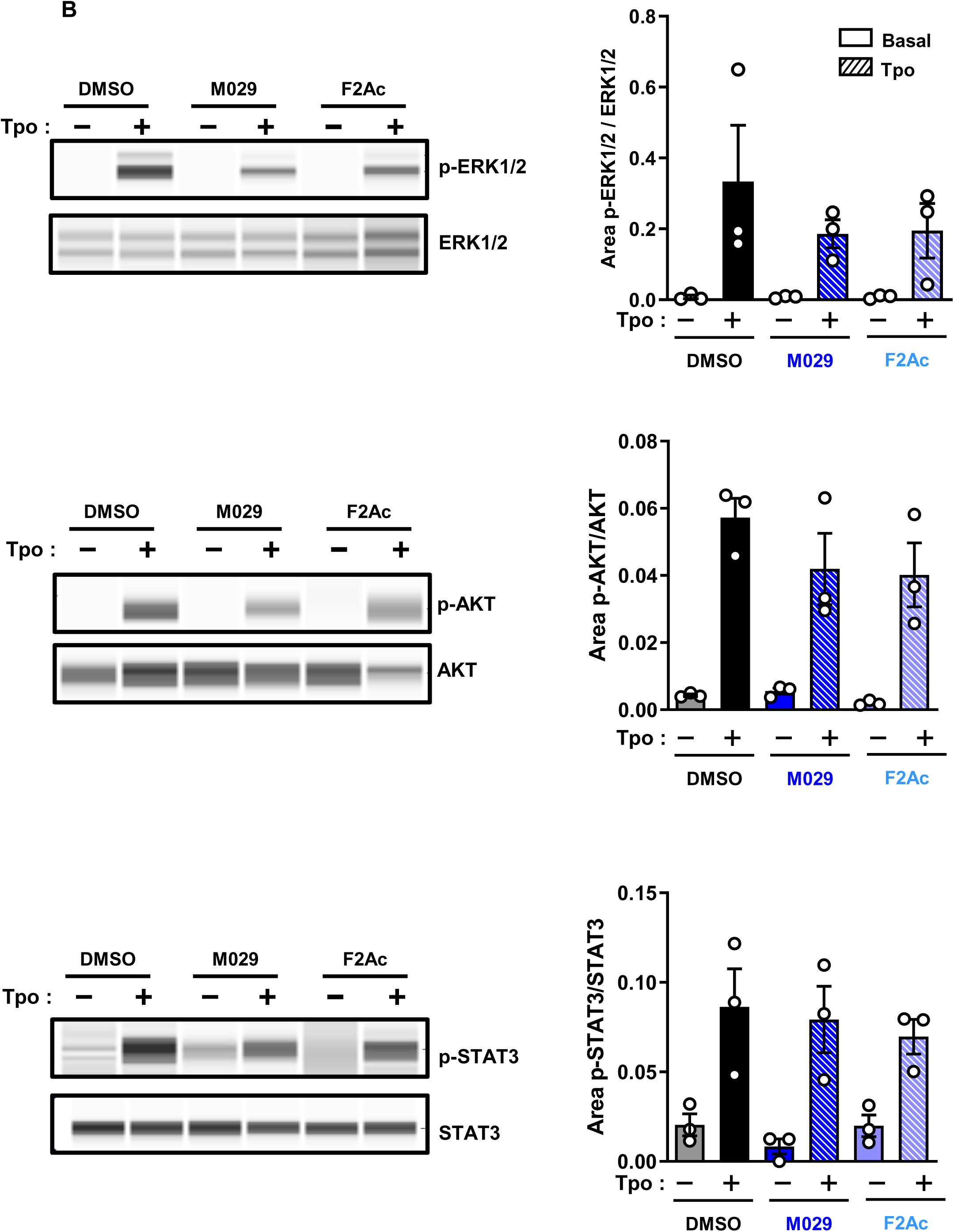
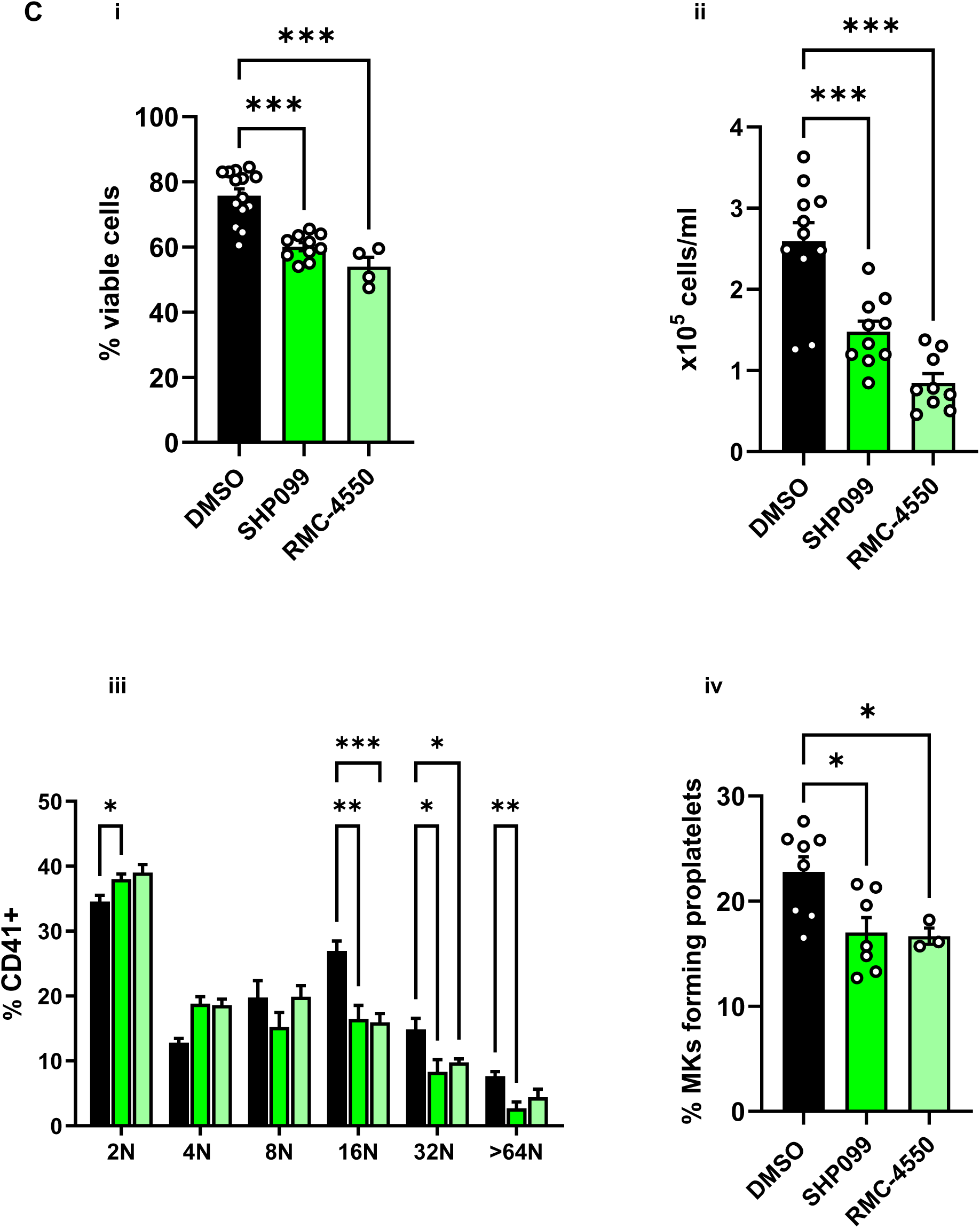
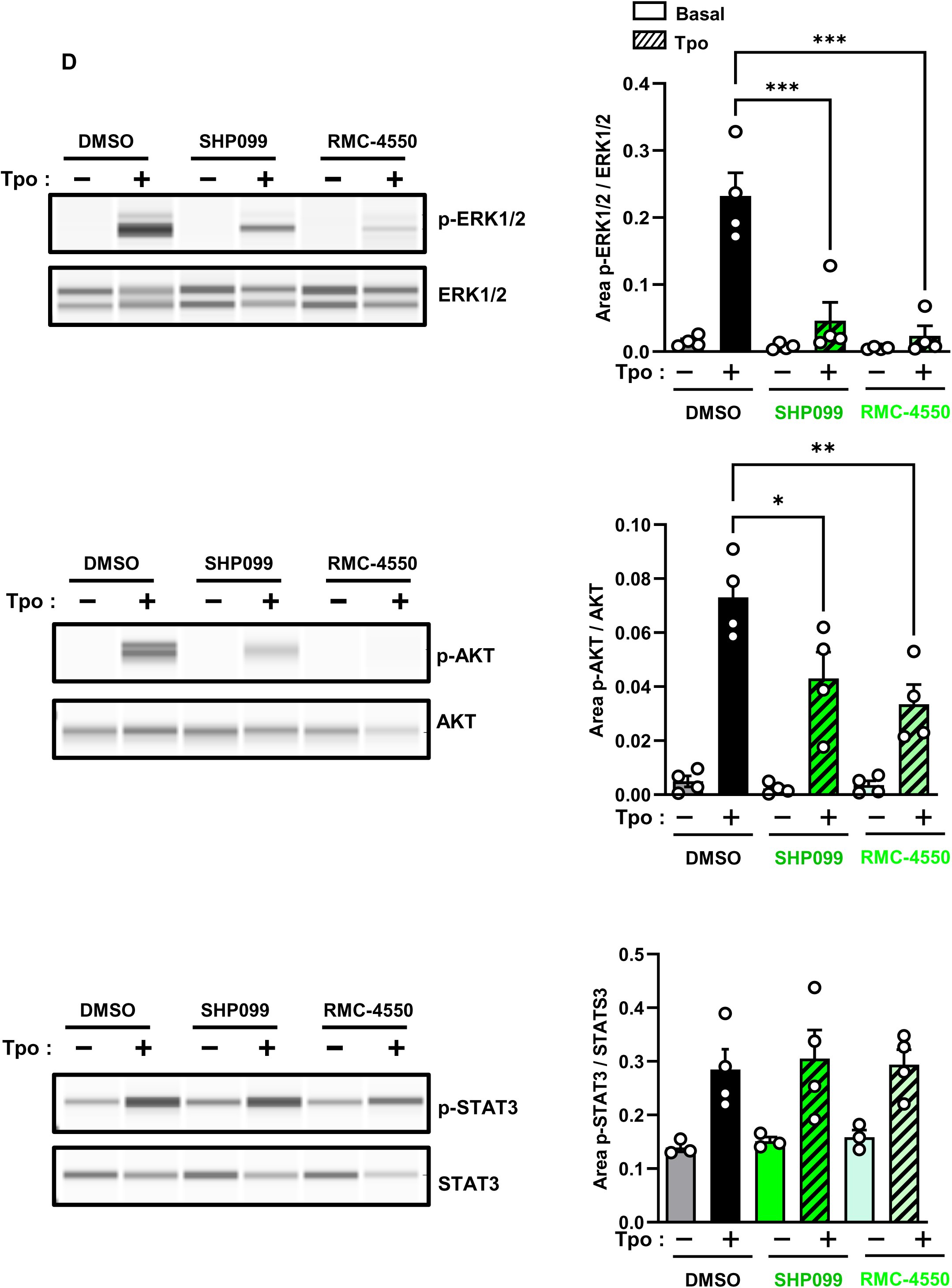
Inhibition of Shp2 activity impairs megakaryopoiesis and Mpl signaling. **(A)** Effects of two selective Shp1 allosteric inhibitors 10 µM M029 and 10 µM F2Ac were tested on megakaryopoeisis. **(i)** Viability, **(ii)** proliferation, and **(iii)** MK ploidy. Quantification from n = 3-4 independent experiments/condition. **(B)** Whole cell lysates of resting and 50 ng/mL Tpo-stimulated MKs were western blotted with the indicated antibodies. Representative blots capillary-based immunoassays and quantification of peak areas from n = 3 independent experiments/genotype. **(C)** Effects of two selective Shp2 allosteric inhibitors 10 µM SHP099 and 10 µM RMC-4550 were tested on megakaryopoeisis. **(i)** Viability, **(ii)** proliferation, **(iii)** MK ploidy and **(iv)** percentage of MKs forming proplatelets. Quantification from n = 3-6 independent experiments/condition. * *P* < 0.05; ** *P* < 0.01; *** *P* < 0.001, two-way ANOVA. **(D)** Whole cell lysates of resting and 50 ng/mL Tpo-stimulated MKs were western blotted with the indicated antibodies. Representative blots capillary-based immunoassays and quantification of peak areas from n = 3 independent experiments/genotype. * *P* < 0.05; ** *P* < 0.01; two-way ANOVA.

## Discussion

In this study, we used the *Gp1ba-Cre* transgenic mouse model to achieve conditional deletion of the tyrosine phosphatases Shp1 and Shp2 in the MK/platelet lineage. Our results demonstrate efficient deletion of Shp1 and Shp2 in MKs and more completely in platelets, confirming the utility of this mouse model for studying protein involvement in megakaryopoiesis and thrombopoiesis. Residual Shp1 and Shp2 in MKs allowed us to study the phenotype of DKO MKs more thoroughly compared with DKO MKs on the *Pf4-Cre* background, which were severely compromised developmentally and did not survive *in vitro*.^15^ Additionally, the less severe MK defects meant fewer platelet anomalies, which contained lower levels of Shp1 and Shp2 than the MKs, suggesting different half-lives of the phosphatases in MKs versus platelets. The *Gp1ba-Cre* transgenic mouse allowed us to reveal distinct and synergistic functions of Shp1 and Shp2 in these processes, without confounding effects of deleting Shp1 and Shp2 in non-MK/platelet lineages observed with the *Pf4-Cre* transgene mouse.

Deletion of Shp1 or Shp2 alone did not significantly affect platelet counts or volumes, while combined deletion resulted in macrothrombocytopenia characterized by a 40% reduction in platelet counts and marginal increase in platelet volume. This phenotype was accompanied by prolonged bleeding and increased blood loss following tail tip excision, indicating impaired hemostatic function. These defects correlated with reduced GPVI and α2 expression, GPVI-mediated platelet aggregation and P-selectin expression. GPVI signaling was also impaired in Shp1/2-deficient platelets, with reduced Syk phosphorylation despite normal SFK activation, suggesting that Shp1 and Shp2 modulate ITAM/Syk proximal signaling (**Figure 8**). This is further supported by the marginal reduction in Syk activation downstream of the hemi-ITAM-containing CLEC-2 receptor. However, the marginal albeit significant reduction in Syk phosphorylation downstream of CLEC-2 in Shp1 and Shp2 KO platelets was not determined and was insufficient to impact CLEC-2-mediated platelet aggregation under the conditions tested. Differences in the stoichiometry and docking of Syk to phosphorylated GPVI-FcR γ-chain and CLEC-2 likely contribute to the differences in platelet reactivity and Syk phosphorylation downstream of the two receptors in the absence of Shp1 and Shp2.

**Figure 8.**
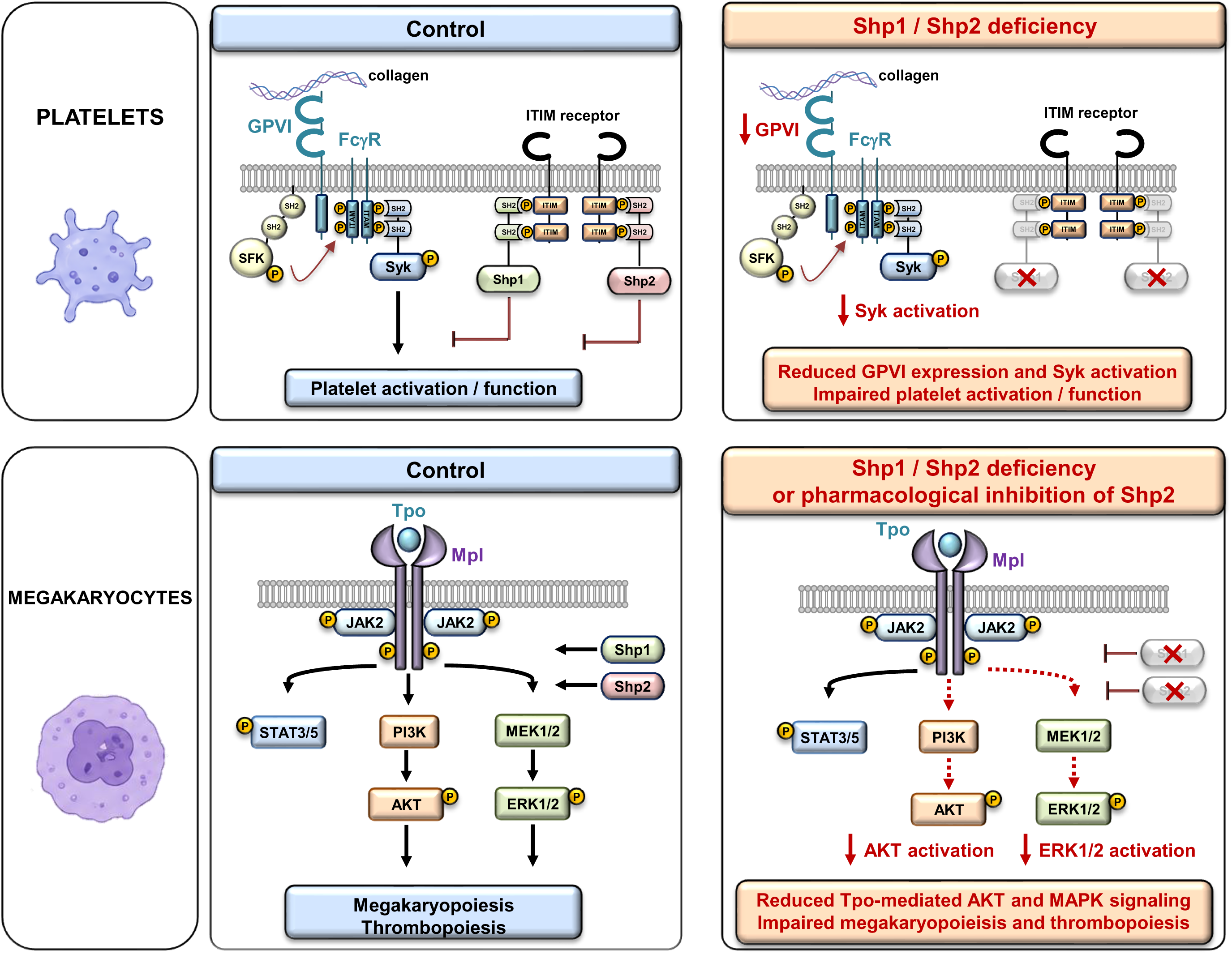
Distinct roles of Shp1 and Shp2 in platelet GPVI signaling and megakaryocyte Tpo/Mpl signaling. Schematic representation summarizing the effects of Shp1/Shp2 deficiency on platelet activation and megakaryopoiesis. In platelets, GPVI–FcRγ engagement by collagen induces ITAM phosphorylation and recruitment of Src family kinases and Syk, leading to platelet activation and aggregation under physiological conditions. Combined Shp1/Shp2 deficiency impairs GPVI expression and Syk activation, resulting in defective platelet activation and aggregation. In megakaryocytes, thrombopoietin (Tpo) binding to Mpl activates JAK2-dependent downstream signaling pathways, including STAT3/5, MAPK/ERK1/2, and PI3K/AKT, thereby promoting normal megakaryopoiesis and platelet production. Shp1/Shp2 deficiency or pharmacological inhibition of Shp2 reduces Tpo-induced ERK1/2 and AKT activation, leading to impaired megakaryocyte differentiation and thrombopoiesis.

Similar phases and kinetics of platelet recovery following immune-induced platelet clearance suggest that thrombocytopenia in Shp1/2 DKO mice is not due to increased platelet clearance, but rather to impaired MK development, maturation and function. *In vitro* assays showed significant defects in MK polyploidization and proplatelet formation, while *in vivo* analysis demonstrated an accumulation of immature MKs with incomplete demarcation membrane system formation and lower ploidy levels. These findings point to a critical requirement for Shp1 and Shp2 in late-stage MK maturation, a process tightly regulated by Tpo signaling through Mpl. Consistently, Tpo-mediated activation of the ERK1/2 pathway was markedly reduced in Shp2- and Shp1/2-deficient MKs, while changes in AKT and STAT3 pathways were modest, underscoring the prominent role of Shp2 in facilitating Tpo-driven Ras/MAPK signaling (**Figure 8**). Based on these findings, the synergistic effects of deleting both Shp1 and Shp2 suggests that Shp1 acts through another, as yet undefined pathway vital for megakaryopoiesis and thrombopoiesis. RhoA/ROCK is an obvious candidate that Shp1 has been implicated in regulating in other lineages and is essential for cytoskeletal remodeling during endomitosis.

Pharmacological inhibition corroborated genetic findings. Selective inhibition of Shp2 with allosteric inhibitors SHP099 and RMC-4550 significantly impaired MK viability, proliferation, polyploidization, proplatelet formation, and Tpo signaling, while Shp1 inhibitors M029 and F2Ac had no significant effects. This demonstrates a dominant role for Shp2 in regulating MK proliferation and differentiation downstream of Tpo, whereas Shp1 is dispensable for these functions. The pharmacological data thus confirm the functional specificity of the two phosphatases and provide further mechanistic insights into their differential contributions. The relatively mild platelet and hemostatic phenotypes observed in Shp1/2 DKO platelets suggest that Shp2 inhibition may have a limited impact on bleeding risk. However, given that these findings are derived from murine models, further studies are required to evaluate the hemostatic consequences of Shp2 inhibition in human systems and in disease-relevant contexts, which is part of a follow-up study.

The relatively milder, lineage-specific phenotype of *Gp1ba-Cre*-driven Shp1/2 DKO mice provides distinct advantages to the previously described *Pf4-Cre* mouse model ^32,33^ **(Supplemental Table 4)**. *Pf4-Cre*-mediated deletion induces recombination earlier during megakaryopoiesis, starting in MK progenitors, whereas *Gp1ba-Cre* is activated predominantly in mature MKs and platelets. This temporal difference allows normal early MK development and platelet production, resulting in less severe thrombocytopenia and functional defects. Additionally, our protein quantification data show significant, but incomplete ablation of Shp1 and Shp2 in BM-derived MKs, suggesting residual phosphatase activity is sufficient under steady-state conditions. In contrast, *Pf4-Cre* models generally achieve near-complete deletion at earlier stages of development, leading to profound thrombocytopenia and often lethality. Furthermore, compensatory extramedullary hematopoiesis observed in our *Gp1ba-Cre* model may further alleviate thrombocytopenia. Importantly, this milder phenotype allowed us to study the Shp1/2 DKO mice in detail, which was impossible with the *Pf4-Cre* model due to the severity of the phenotype and cell viability. Findings further establish the utility of the *Gp1ba-Cre* strain for dissecting the distinct roles of Shp1 and Shp2 in late-stage megakaryopoiesis and platelet biology.

Collectively, Shp2 is a critical positive regulator of Mpl signaling and downstream Ras/MAPK pathways, essential for MK proliferation, polyploidization, and proplatelet formation, whereas Shp1 plays a more modulatory role in ITAM-containing receptor signaling in platelets without directly impacting MK maturation. The combined deletion of Shp1 and Shp2 reveals additive effects on platelet function and MK development, underscoring the importance of coordinated phosphatase regulation in maintaining platelet homeostasis. This study provides the first detailed analysis of Shp1 and Shp2 functions in megakaryopoiesis and platelet biology using a late-stage MK/platelet-specific deletion model, highlighting the indispensable role of Shp2 in megakaryopoiesis and thrombopoiesis, and distinct contributions of Shp1 in platelet signaling. It also emphasizes the utility of the *Gp1ba-Cre* mouse for exploring protein function in the MK/platelet lineage. These insights may have broader implications for understanding phosphatase regulation in hematopoiesis and could inform therapeutic targeting of Shp2 in hematologic diseases.

## Acknowledgements

This work was supported by Inserm, the Agence Nationale pour la Recherche (AM, ANR-22-REMYS-CE14-0061-01 (E. Barre, PhD candidate), and YS, ANR-23-TARGITT-CE18-0020-01) and the European Research Council (YS ERC-MENTOR-101141783. ZQ, JM and ZYZ are supported by NIH RO1 CA069202. The authors acknowledge all members of the animal facility at UMR_S1255 for maintenance of mouse colonies and the flow cytometry platform (CytoTriCS).

## Author Contributions

E.B. Performed experiments, analyzed data, revised the manuscript.

MD.LR. Performed experiments, analyzed data, revised the manuscript.

L.Z. Performed experiments and analyzed data.

M.P. Performed experiments and analyzed data.

C.L. Performed experiments and analyzed data.

F.P. Performed experiments and analyzed data.

JY.R. Performed experiments and analyzed data.

A.E. Analyzed data.

Z.Q. Provided reagents.

J.M. Provided reagents.

Z.Y.Z. Provided reagents and revised the manuscript.

Y.A.S. Designed experiments, analyzed data, wrote and revised the manuscript.

A.M. Conceptualized, designed experiments, analyzed data, wrote and revised the manuscript.

## Data Availability Statement

The data that support the findings of this study are available from the corresponding author upon reasonable request.

