## Supplemental material for "Synergistic effects of deleting the tyrosine phosphatases Shp1 and Shp2 on megakaryopoiesis and thrombopoiesis in mice"

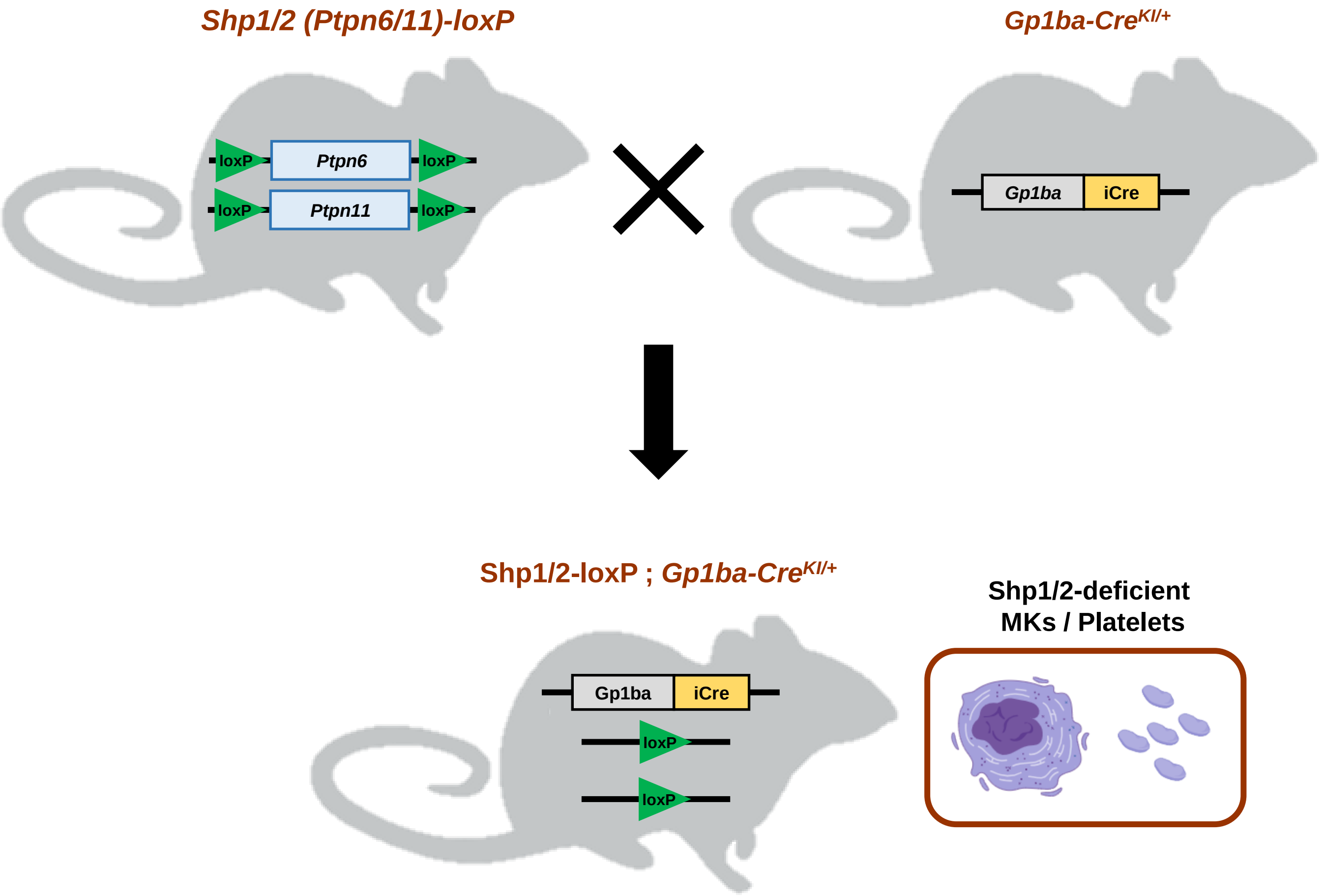


**Figure S1. Breeding schemes of the generation of *Ptpn6/Ptpn11*$^{flox/flox}$ ; *Gp1ba-Cre*$^{KI/+}$ mice.** MK/Platelet specific Shp1 and Shp2 knockout (KO) mice were generated by crossing *Ptpn6*$^{fl/fl}$ and *Ptpn11*$^{fl/fl}$ mice (Shp1/2-loxP) with the *Gp1ba-Cre* transgenic deleter mouse *Gp1ba-Cre*$^{KI/+}$.

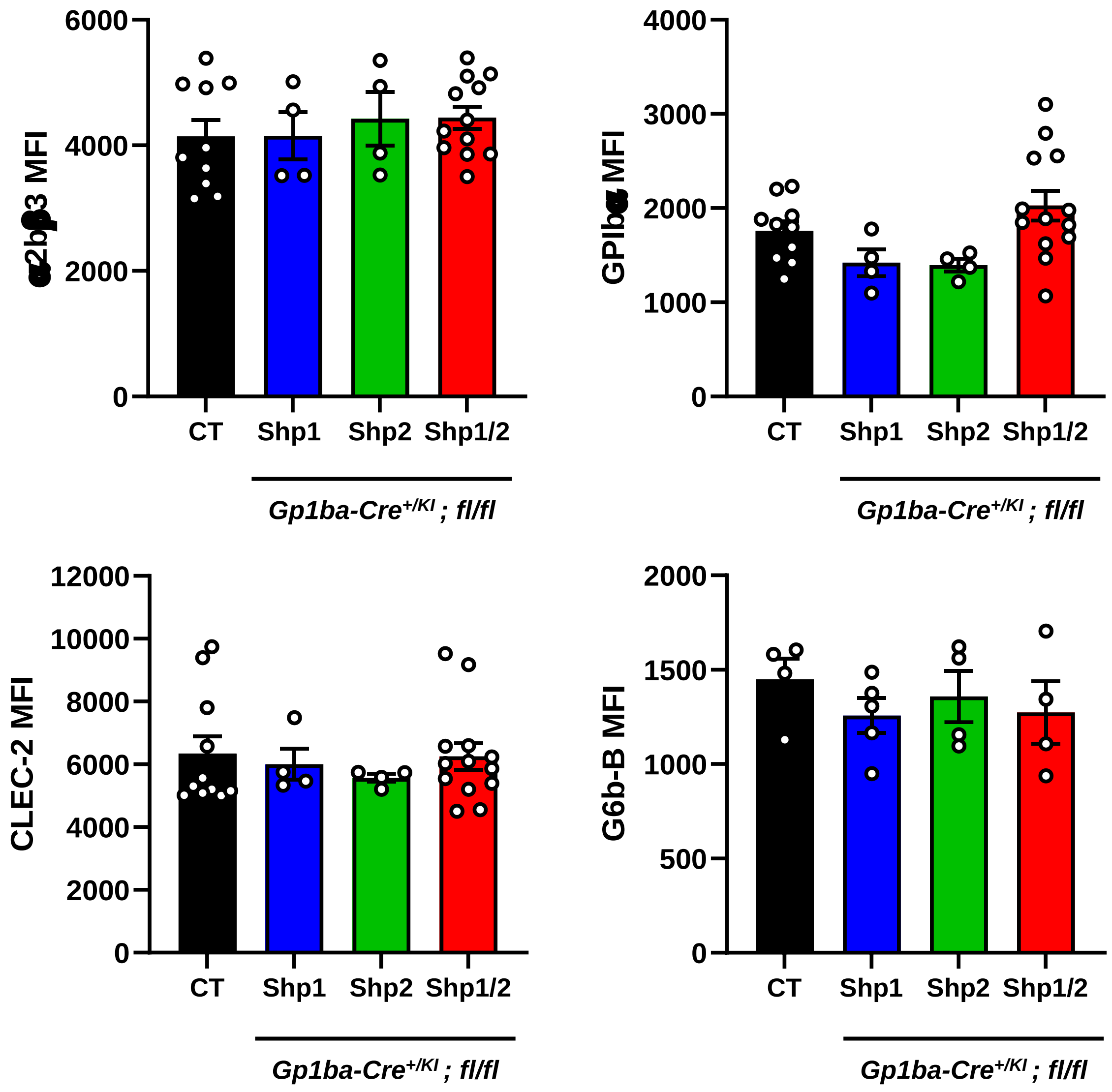


**Figure S2. Surface glycoprotein expression.** Platelet surface receptor expression of integrin αIIbβ3, GPIbα, CLEC-2 and G6b-B were measured in whole blood by flow cytometry and shown as median fluorescence intensity (MFI), n = 4 to 10 mice per genotype.

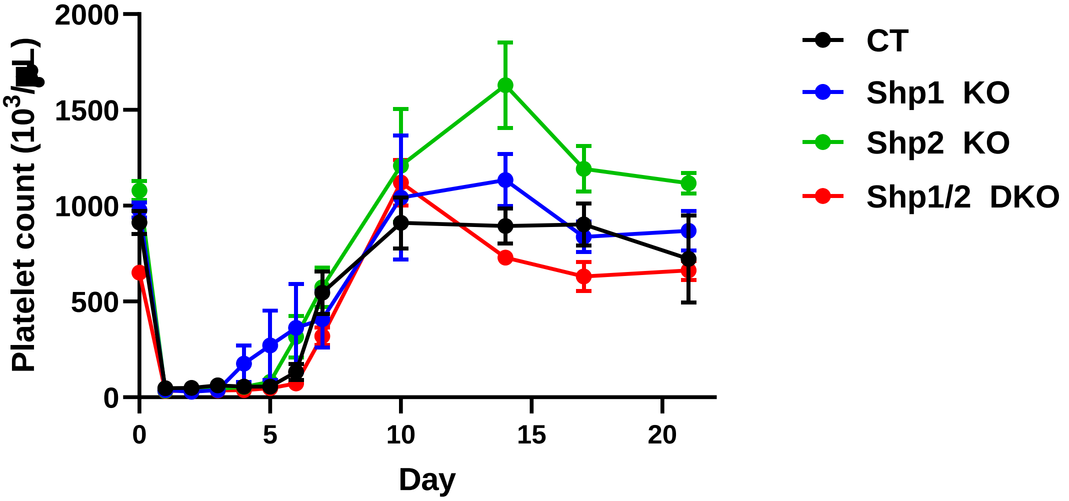


**Figure S3. Platelet recovery following antibody-mediated depletion.** Platelet counts were measured prior and following anti-GPIbα antibody-mediated platelet depletion at indicated time points (n = 3-8 mice/time point/genotype; mean ± SEM). No statistically significant differences between genotypes were observed using two-way ANOVA analysis.

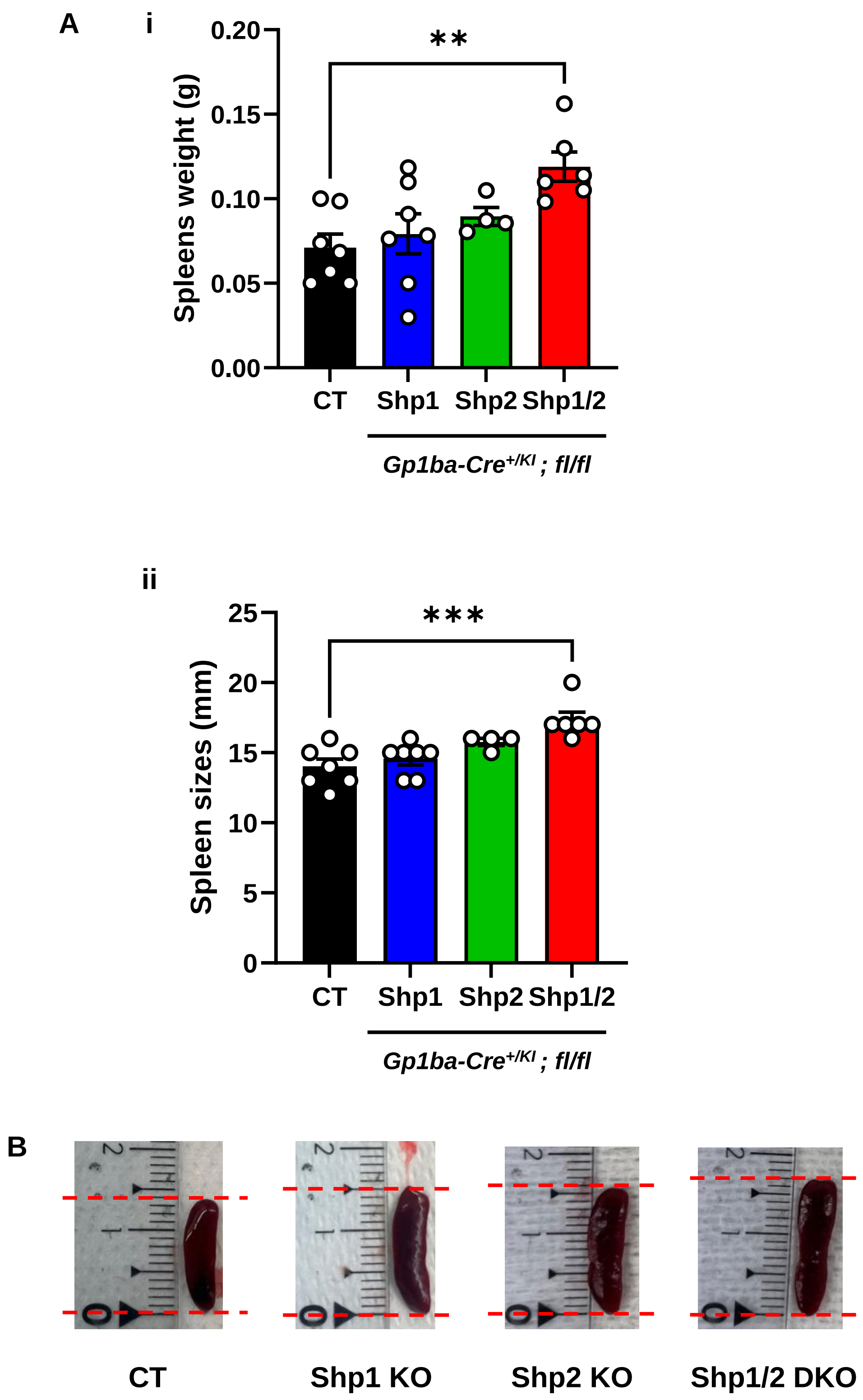


**Figure S4. Splenomegaly in Shp1/2 DKO mice. (A)** Significant increases in spleen **(i)** weight and **(ii)** size in Shp1/2 DKO mice (n = 7). **(B)** Representative images. Mean ± SEM, one-way ANOVA, ; ** $p < 0.01$ ; *** $p < 0.001$.

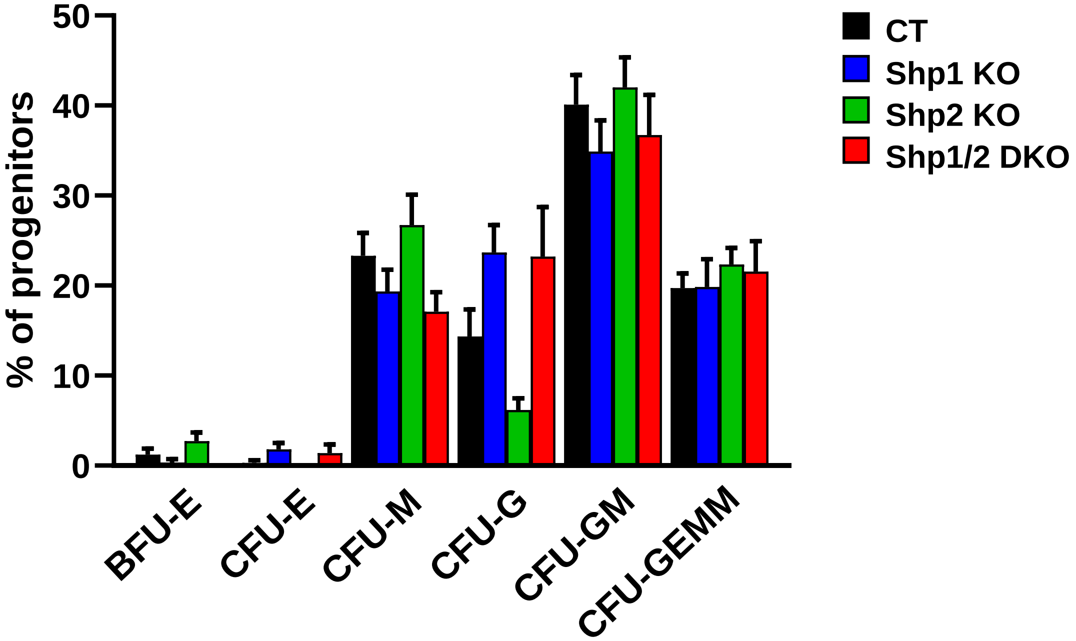


**Figure S5. Colony-forming unit (CFU) assay of hematopoietic stem/progenitor cells.** Mouse HSPCs were isolated from bone marrow and plated in methylcellulose-based medium containing appropriate cytokines (MethoCult™). Cells were cultured for 10–14 days at 37 °C with 5% $CO_2$. Colonies were identified, counted and classified into BFU-E (burst-forming unit-erythroid), CFU-E (erythrocyte), CFU-M (macrophage), CFU-G (granulocyte), CFU-GM (granulocyte-macrophage) and CFU-GEMM (granulocyte, erythrocyte, macrophage, megakaryocyte). Bar graphs represent the mean ± SEM of n = 8 -14 independent experiments. Statistical significance was determined using two-way ANOVA.

**Table 1. Antibodies and working concentrations for JESS experiments.**

| Antibody | References | Dilution | Protein concentration (MKs) | Protein concentration (Platelets) |
|---|---|---|---|---|
| SHP-1 | 3759 – Cell Signaling | 1/10 | 0.2 mg/ml | 0.2 mg/ml |
| SHP-2 | Sc-7384 – Santa Cruz | 1/10 | 0.2 mg/ml | 0.2 mg/ml |
| ERK1/2 | 9102 – Cell Signaling | 1/25 | 0.2 mg/ml | 0.2 mg/ml |
| p-ERK1/2 (Thr202/Tyr204) | 9101 – Cell Signaling | 1/25 | 0.2 mg/ml | - |
| STAT3 | 4904 – Cell Signaling | 1/50 | 0.2 mg/ml | - |
| p-STAT3 (Tyr705) | 9145 – Cell Signaling | 1/10 | 0.2 mg/ml | - |
| AKT | 4691 – Cell Signaling | 1/50 | 0.2 mg/ml | - |
| p-AKT (Ser473) | 9271 – Cell Signaling | 1/20 | 0.2 mg/ml | - |
| Syk p-Tyr519/520 | 2711 – Cell Signaling | 1/50 | - | 0.1 mg/ml |
| Syk | 12358 – Cell Signaling | 1/50 | - | 0.1 mg/ml |
| Src p-Tyr418 | 44660G – Invitrogen | 1/25 | - | 0.1 mg/ml |
| Src | 2108 – Cell Signaling | 1/25 | - | 0.1 mg/ml |

**Table 2. Hematological analysis of control mice (*Gp1ba-Cre* $^{KI/+}$), Shp1 KO, Shp2 KO and Shp1/2 DKO mice.**

| Parameters | Gp1ba-Cre$^{KI/+}$ (n = 79) | Shp1 KO (n = 26) | Shp2 KO (n = 39) | Shp1/2 DKO (n = 53) |
|---|---|---|---|---|
| PLT ($10^3$/µL) | 827.04 ± 205.8 | 867.69 ± 203.2 | 963.28 ± 247.7 | 631.66*** ± 258.8 |
| MPV (flt) | 5.94 ± 0.43 | 5.75 ± 0.42 | 6.25 ± 0.54 | 6.93*** ± 0.85 |
| RBC ($10^6$/µL) | 7.43 ± 1.23 | 8.48 ± 1.15 | 7.76 ± 1.03 | 7.65 ± 1.16 |
| HCT (%) | 34.64 ± 6.01 | 39.55 ± 5.60 | 36.29 ± 5.10 | 36.63 ± 5.57 |
| WBC ($10^3$/µL) | 10.44 ± 4.04 | 9.60 ± 4.05 | 12.29 ± 5.99 | 11.94 ± 8.39 |
| LYM ($10^3$/µL) | 7.93 ± 3.46 | 6.89 ± 3.02 | 7.63 ± 4.88 | 6.71 ± 3.78 |
| MON ($10^3$/µL) | 0.36 ± 0.28 | 0.56 ± 0.76 | 1.45 ± 1.73 | 1.82*** ± 3.17 |
| NEU ($10^3$/µL) | 2.01 ± 1.50 | 1.94 ± 1.04 | 2.82 ± 1.88 | 3.02* ± 3.32 |
| EOS ($10^3$/ml) | 0.14 ± 0.11 | 0.20 ± 0.23 | 0.37 ± 0.29 | 0.37*** ± 0.74 |
| BAS ($10^3$/µL) | 0.00 ± 0.01 | 0.00 ± 0.01 | 0.01 ± 0.02 | 0.03* ± 0.07 |

PLT, platelets ; MPV , mean platelet volume; RBC, red blood cells ; HCT, haematocrit, WBC, white blood cells ; LYM, lymphocytes ; MON, monocytes ; NEU, neutrophils ; EOS, eosinophils ; BAS, basophils ; Mean ± standard error ,* $P < 0.05$,*** $P < 0.001$,one-way ANOVA with Sidak's test.

**Table 3. Platelet surface glycoprotein expression in whole blood in control mice (*Gp1ba-Cre* $^{KI/+}$), Shp1 KO, Shp2 KO and Shp1/2 DKO mice.**

| Surface glycoproteins | Gp1ba-Cre$^{KI/+}$ (n = 10) | Shp1 KO (n = 5) | Shp2 KO (n = 5) | Shp1/2 DKO (n = 14) |
|---|---|---|---|---|
| GPVI | 1681 ± 48.14 | 1344.50 ± 170.60 | 1320.25 ± 140.12 | 1078.69 ± 34.68 *** |
| Integrin α2 | 1102.90 ± 43.61 | 1064.50 ± 109.72 | 1083 ± 79.28 | 874 ± 33.13 *** |
| Integrin α2bβ3 | 4140.80 ± 254.76 | 4153.50 ± 434.92 | 4423.25 ± 497.96 | 4440.67 ± 163.92 |
| GPIbα | 1758.20 ± 98.30 | 1420 ± 164 | 1395 ± 76.73 | 2027.31 ± 152.04 |
| CLEC-2 | 6327.70 ± 573.91 | 6007.75 ± 577.48 | 5567.25 ± 143.67 | 6251.15 ± 408.16 |
| G6b-B | 1555.67 ± 19.78 | 1257 ± 119 | 1358.50 ± 156.57 | 1273.25 ± 88.76 |

**Table 4. Mouse Shp1/2 conditional knockout phenotypes on *Gp1ba-Cre* and *Pf4-Cre* backgrounds.**

| | | Genotype / Phenotype | | | | | |
|---|---|---|---|---|---|---|---|
| | | *Pf4-Cre*$^{+}$ | | | *Gp1ba-Cre*$^{KI/+}$ | | |
| **Cell type** | **Parameter** | **Shp1 KO** | **Shp2 KO** | **DKO** | **Shp1 KO** | **Shp2 KO** | **DKO** |
| **MK** | **Ploidy** | ↑ ++ 2-4N<br>↓ ++ 8-128N | ↑ + 2-4N<br>↓ ++ 8-128N | ↑ ++ 2-4N<br>↓ ++ 8-128N | | | ↑ + 2-8N<br>↓ ++ 16-64N |
| | **Function** | | ↓ spreading<br>↑ Tpo level | ↑ + Tpo level | | | ↓ ++ PPT<br>↑ ++ MK stage II<br>↓++ MK stage III |
| | **Signaling** | | ↓ p-ERK1/2 | +++<br>↓ p-ERK1/2 | | ↓ ++<br>p-ERK1/2 | ↓ ++<br>p-ERK1/2 |
| **Platelets** | **Count** | | ↓ ++ | ↓ ++ | | | ↓ ++ |
| | **Volume** | ↑ + | ↑ ++ | ↑ ++ | | | ↑ ++ |
| | **Function** | ↓ CRP aggregation<br>↓ + spreading on fibrinogen<br>↓ + activation (P-selectin) | ↑ CLEC2 aggregation<br>↑ spreading on fibrinogen<br>↓ half-life and platelet recovery | ↓ + spreading on fibrinogen<br>↓ activation (P-selectin)<br>↓ half-life | | | ↑ blood loss<br>↓ CRP and collagen aggregation<br>↓ ++ activation (P-selectin) |
| | **Receptors** | ↓ GPVI | | ↓ +++ | | | ↓ ++ GPVI and α2 |
| | **Signaling** | ↓ p-Src<br>↓ p-Syk | | ↓ +++ | | | ↓ ++ p-Syk |

Blank cell : no defect

+, ++, +++ : mild, medium and severe defect

Shp1 KO : *Pf4-Cre*$^{+}$ ; *Ptpn6*$^{fl/fl}$ or *Gp1ba-Cre*$^{KI/+}$ ; *Ptpn6*$^{fl/fl}$

Shp2 KO : *Pf4-Cre*$^{+}$ ; *Ptpn11*$^{fl/fl}$ or *Gp1ba-Cre*$^{KI/}$ ; *Ptpn11*$^{fl/fl}$

Shp1/2 DKO : *Pf4-Cre*$^{+}$ ; *Ptpn6*$^{fl/fl}$ ; *Ptpn11*$^{fl/fl}$ or *Gp1ba-Cre*$^{KI/+}$ ; *Ptpn6*$^{fl/fl}$ ; *Ptpn11*$^{fl/fl}$

MK : megakaryocytes

PPT : proplatelet megakaryocytes
